# ONETest PathoGenome: A Multi-Cohort Evaluation of an Optimized NGS Assay for Detection of Lower Respiratory Pathogens in Bronchoalveolar Lavage

**DOI:** 10.64898/2026.03.26.714510

**Authors:** Sepideh Massoumi Alamouti, Hoang Dong Nguyen, Habib Daneshpajouh, Noushin Moshgabadi, Brian S Kwok, Herbert J Houck, Greg Stazyk, Tang Patrick, Cherabuddi Kartikeya, Petr Starostik, Mohammad A. Qadir, Kenneth H Rand

## Abstract

**Background:** Lower respiratory tract infections (LRTIs) remain diagnostically challenging when culture and molecular assays are negative or delayed. We evaluated ONETest™ Pathogenome (OT), an automated hybrid-capture metagenomic assay with core-genome enrichment probes, for direct pathogen detection in bronchoalveolar lavage (BAL).

**Methods:** Analytical performance (LoD, precision, continuity) was assessed using whole-cell spike-ins into culture-negative BAL fluid. Technical performance was assessed in 119 specimens profiled by OT and whole-metagenome shotgun sequencing (WmGS, cohort 1). Clinical accuracy was evaluated in 360 specimens (cohort 2) benchmarked against routine bacterial and acid-fast bacillus (AFB) culture. Laboratory-developed test (LDT) validation included 43 specimens (cohort 3) benchmarked to bacterial and AFB culture.

**Results:** OT uses 6.2 million probes covering core genomes across 50 microbial families (>250 respiratory pathogens). In BAL specimens, OT increased normalized on-target microbial abundance 26-fold versus that of WmGS while preserving within-sample microbial diversity. In cohort 2, OT achieved species-level sensitivity of 80% and specificity of 99% across culture-confirmed isolates and detected ≥1 culture-confirmed organism in 100/115 culture-positive specimens (87%), while applying species-specific background baselines to mitigate overcalling. Additive yield was 21% (76/360), with 7.5% (27/360) of specimens having ≥1 additional finding supported by orthogonal testing. In LDT validation, OT identified ≥1 culture-confirmed organism in 34/40 culture-positive specimens (85%) with one OT-positive/culture-negative specimen.

**Conclusions:** OT is an assay with a turnaround time <24 h complementary to culture that improves pathogen detection and expands microbiologic findings through additional detections and co-detections, including slow-growing organisms that may require prolonged incubation by conventional methods.

## INTRODUCTION

LRTIs remain a leading cause of mortality among hospitalized patients worldwide^1^ requiring timely etiologic diagnosis to enable targeted therapies. Current care standards rely on a combination of clinical evaluation, imaging, and laboratory diagnostics, including Polymerase Chain Reaction (PCR) and cultures^2^. Culture-based methods, while comprehensive, are struggling with slow-growing or non-culturable organisms, and their sensitivity decreases after antibiotic exposure. PCR, on the other hand, offers faster results but remains constrained by predefined multiplex panels, limiting coverage of the full etiologic spectrum^3^.

WmGS has promised a broad array of organism detection within a single test, surpassing culture and PCR in scope^4^. However, its application in clinical microbiology is hampered by cost, labor, and computational demands^5^. A key limitation is that most sequencing reads derive from the host genome, and differentiating causative organism(s) from commensals or contaminants remains challenging, limiting the ability to generate actionable results^5,6^. Additional barriers include assay complexity and lack of automation^5^.

Targeted metagenomic sequencing (TmGS) is an alternative to WmGS that improves efficiency^5^ by focusing on specific genomic regions using PCR-based amplicon enrichment or probe-based hybrid capture reducing sequencing burden and improving signal-to-noise ratios^5^. Amplicon-based methods such as 16S rRNA sequencing typically offer genus-level resolution and cannot reliably distinguish closely related species^7^, while probe-based assays are constrained by the need to anticipate clinically relevant pathogens and may miss novel or divergent strains^5^. To overcome these limitations while retaining broad coverage, we developed OT (Fusion Genomics Corp. Richmond, BC, Canada), a TmGS assay that captures thousands of core-genome loci across 254 microbial species and their genotypes enabling detection of major pathogens implicated in LRTIs. OT offers a fully automated DNA to results workflow with integrated bioinformatic analysis. Here, we assess the analytical performance of a 6-million-QuantumProbe (QP) enrichment panel by determining the limit of detection (LoD) in BAL fluid spiked with cultured pathogen cells, compare OT to WmGS in 119 BAL specimens, and evaluate diagnostic performance in 360 specimens from a clinical trial cohort benchmarked against culture. Finally, we report LDT validation results from 43 BAL specimens to support regulatory and clinical implementation.

## METHODS

### Study population

The study was conducted at the University of Florida (UF) Health Shands Hospital in Gainesville, Florida, a major tertiary care center (>1,100 beds) with a large transplant population serving North Central Florida. The study included 537 patients across three cohorts: (1) 119, (2) 375, and (3) 43 samples. All patients underwent BAL as part of routine standard care between 2022 – 2024, with specimens submitted for cell count, culture, and molecular testing (Fig. S2, Table S2). For the clinical trial population (n=360, cohort 2) consecutive BAL specimens were collected between 12/6/2022 and 11/13/2023 and stored frozen at -70°C after performance of the cell differential in the Hematology laboratory until they could be further studied. The mean age was 54.1 ± 22.5 years, range 0 – 90, 212 (58.9%) were male, and 142 had received organ transplantation, primarily lung (N=112). Demographic data and reasons for admission were available only for the cohort 2. Exclusion criteria included quantity not sufficient (QNS); in cohort 2, 15/375 specimens were excluded from the final analysis based on an internal quality assessment using background human-read levels described in supplementary method. Additional conventional testing was performed, as directed by the ordering physician according to clinical indications including acid-fast bacilli (AFB) culture for *Mycobacterium* spp. (n = 315 cohort 2, n=42 cohort 3), BioFire® Pneumonia Panel (BioFire Diagnostics, UT, USA; n = 92), and MGDx testing (Lubbock, TX, USA; n = 66).

### Assay design

OT is an end-to-end, walk-away automated assay that generates enriched WmG libraries for core genomes of LRT pathogens and related lineages (Fig. S1). It is compatible with Illumina sequencing and incorporates data analytics to determine the genomic profile of microorganisms present in a given sample. The analysis applies a multi-parameter score to distinguish sample-derived microbial signal over background and potential contamination. The OT’s current iteration enriches nucleic acids from known pathogens associated with LRTIs and related lineages while also profiling commensal microbiota present in each specimen. The assay includes all reagents required to generate Illumina-compatible NGS libraries from total nucleic acids, and sequencing data are analyzed through the FusionCloud portal. OT also uses multiple in-house custom-reference datasets that support standardized QP design and downstream analysis, as described below.

#### *Microbial qualification and categorization* (summary)

Reported microbial signals were qualified and categorized using a proprietary statistical framework integrated within the OT pipeline (see supplementary method). Classification is informed by empirically derived, species-specific thresholds calibrated on historical clinical BAL samples and dynamically applied to each test, allowing thresholds to be defined based on observed distributions multiple metrics, and classification performance was benchmarked against conventional culture in validation.

#### Analytical performance (LoD)

A reference panel of six clinically relevant organisms was obtained from Shands Hospital at 0.5 McFarland (Table 1). The panel included five bacterial and one fungal species spanning 32–67% GC content and genome sizes of 2–36 Mb and was pooled with BAL samples confirmed negative by culture and OT. The pooled material was serially diluted 10-fold in negative BAL seven times, generating concentrations ranging from 23,000,000 to 23 CFU/mL. Each dilution was prepared in triplicate and independently extracted and sequenced according to the protocol described below.

**Table 1.**
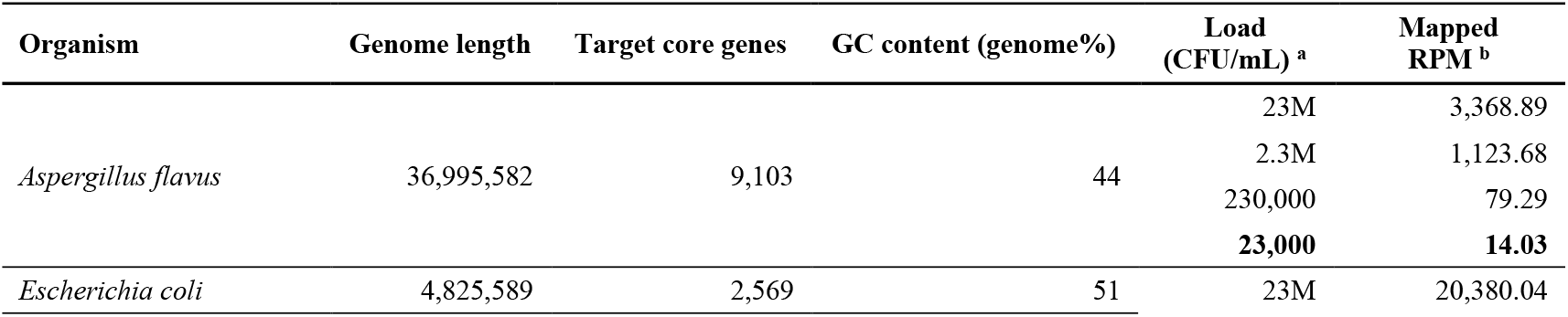
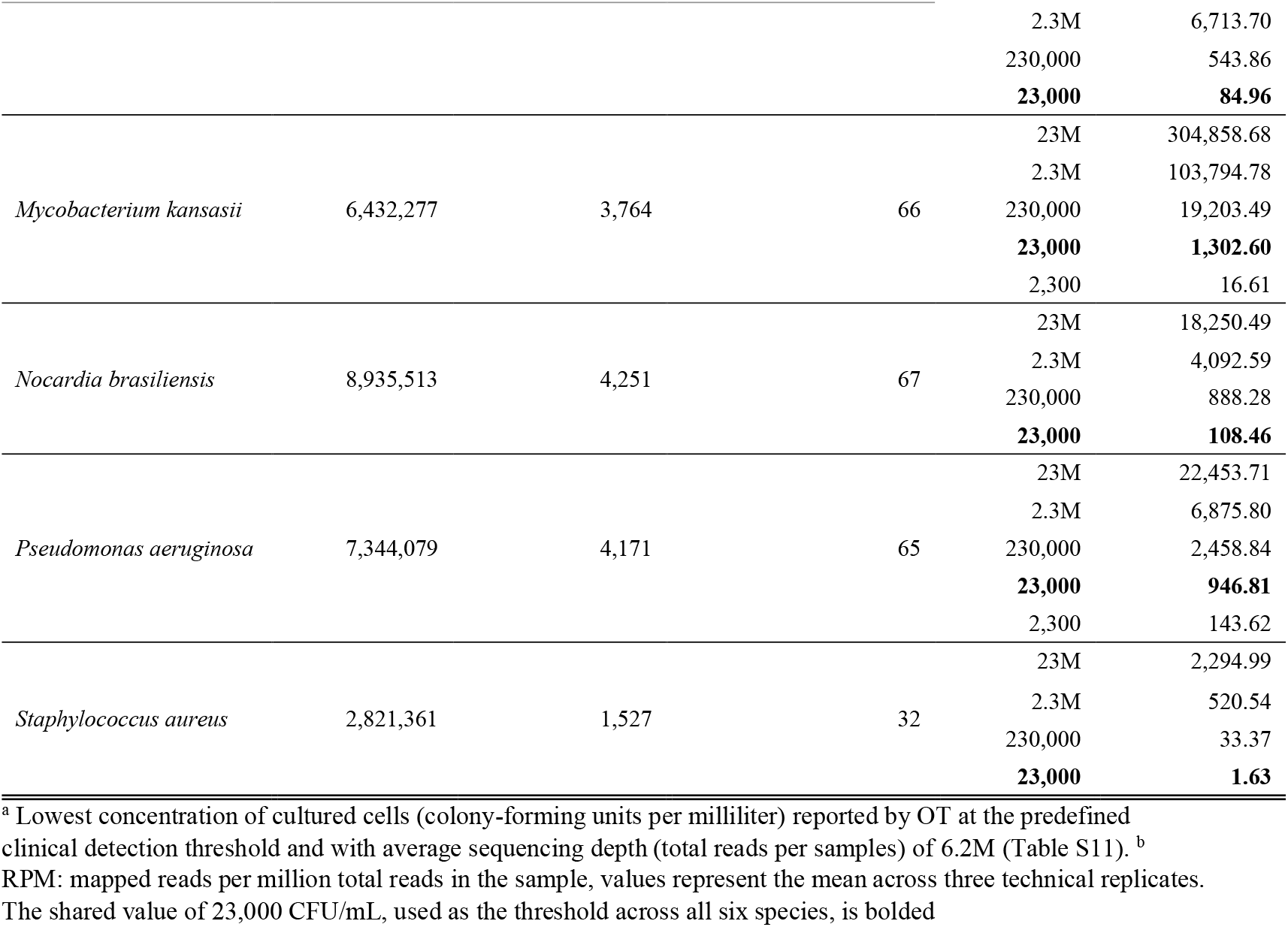
ONETest™ level of detection for microbial cells spiked into BAL clinical specimens.

#### Ethical approval

This study was approved by the UF Institutional Review Board (IRB202202639). Samples were de-identified prior to testing and the study was exempt from requiring individual informed consent

#### Performance and Quality control description

WmGS and OT assays were performed on cohort 1 together with routine culture, whereas only OT and routine culture were performed on cohorts 2 and 3. Cohorts 1 and 2 were processed in batches on 96-well plates containing 84 clinical specimens and 12 controls. Cohort 3 samples, generated for LDT validation, were processed on a Hamilton STAR automated liquid-handling platform in batches of seven clinical specimens plus one negative control per run. For cohorts 1 and 2, controls included positive, negative, and no-template controls (NTC). For cohort 3, controls included positive and negative controls only. The negative control (HCC) consisted of human cell-line genomic DNA (Jurkat; Thermo Scientific, Mississauga, ON, Canada). Positive controls comprised assay reference material prepared from the ZymoBIOMICS™ Microbial Community DNA Standard (Zymo Research, Irvine, CA, USA). An Illumina PhiX control library (1–2% spike-in) was included in each sequencing run in accordance with manufacturer recommendations.

#### Statistical analysis

Conventional diagnostic culture (routine bacterial and AFB culture) served as the reference standard. OT performance was evaluated by estimating sensitivity, specificity, and percent agreement at both the species (per-pathogen) and per-sample levels. Agreement between OT and culture was summarized using Cohen’s κ statistic with 95% asymptotic confidence intervals (R package v1.4.13) and interpreted according to McHugh’s criteria (2012). Within-sample species richness and evenness were quantified using the Shannon diversity index, calculated from species-level relative abundances estimated by Kraken2–Bracken (Kraken2 --confidence 0.3; Bracken -r 150 -l S). Statistical analyses were performed in R version 4.5.1. Group comparisons were conducted using the Wilcoxon rank-sum test, with *p*-values adjusted for multiple comparisons using the Benjamini–Hochberg procedure to control the false discovery rate.

## RESULTS

### Limit of detection (Analytical performance)

To assess the analytical sensitivity of the 6M-QP™ enrichment panel, we measured LoD using whole-cell spike-ins of five bacterial species (*Escherichia coli, Mycobacterium kansasii, Nocardia brasiliensis, Pseudomonas aeruginosa, and Staphylococcus aureus*) and one fungal species (*Aspergillus flavus*) in negative BAL (Table 1). LoD was defined operationally using the assay’s clinical reporting threshold applied in this study. A dilution was considered positive only if the organism was detected in all three technical replicates (3/3) at an average sequencing depth of 6.2M reads per library. Under these conditions, all six species were reported at ≥23,000 CFU/mL (3/3 replicates), while *M. kansasii* and *P. aeruginosa* at 2,300 CFU/mL.

### Precision

Inter-assay reproducibility was assessed by OT testing of three BAL positive samples across three consecutive robot runs, and intra-assay reproducibility by testing three independently prepared sets of these samples within a single run. Internal spiked phage controls met sequencing quality-control criteria in every run, with an average of 4.9M reads per sample, and all sequencing libraries exceeded minimum cutoff of 0.5M reads. OT showed 100% reproducibility, with all five organisms present in positive samples detected at the pre-established clinical detection thresholds across the intra-assay run and each inter-assay run (Table S3).

### Continuity

Continuity measurements were generated for all designated negative-control samples processed alongside clinical BAL specimens in cohort 3. For each LDT run, the Drift Ratio (FPKM *L. monocytogenes* / FPKM human) was evaluated against pre-established acceptance limits. Across the LDT study period, Drift Ratios for negative controls remained within these limits, with no evidence of systematic upward or downward trends, and no runs included in clinical performance analyses were excluded due to continuity control failure. A representative continuity measurement report from a clinical run is shown in Fig. S1.

### Technical performance (cohort 1)

We evaluated the technical efficiency of OT versus WmGS using 119 BAL specimens (cohort 1, Table S2). Across samples, OT and WmGS showed strong agreement in within-sample diversity, with correlated species richness (Fig. 1A; R=0.74; p<0.001) and similar Shannon index values (Table S4). At matched library sizes (Table S5), OT showed a 4.2% absolute reduction in the human-read fraction (p<0.001; Fig. 1B) with increased on-target signal (Fig. 1C), including higher FPKM (mean 26-fold; 95% CI ±2.45; p<0.001) and broader genome coverage (mean 2.42-fold; 95% CI ±0.10; p<0.001). Using full-depth WmGS (mean 28M reads) versus OT (mean 6.9M reads) in the same specimens, OT yielded higher per-pathogen signal for most culture-confirmed species (Fig. 1D, Table S5), with notable enrichments for *P. aeruginosa* (61-fold), *S. pneumoniae* (56-fold), and *S. marcescens* (48-fold). In contrast, OT showed lower signal for non-targeted species (mean 0.48-fold vs WmGS). Finally, among the 28 culture-positive samples, one specimen with strong *M. kansasii* growth was detected by OT but missed by WmGS (Table S6); all other discordances involved specimens annotated as sparse growth or cultures dominated by a different organism.

**Figure 1:**
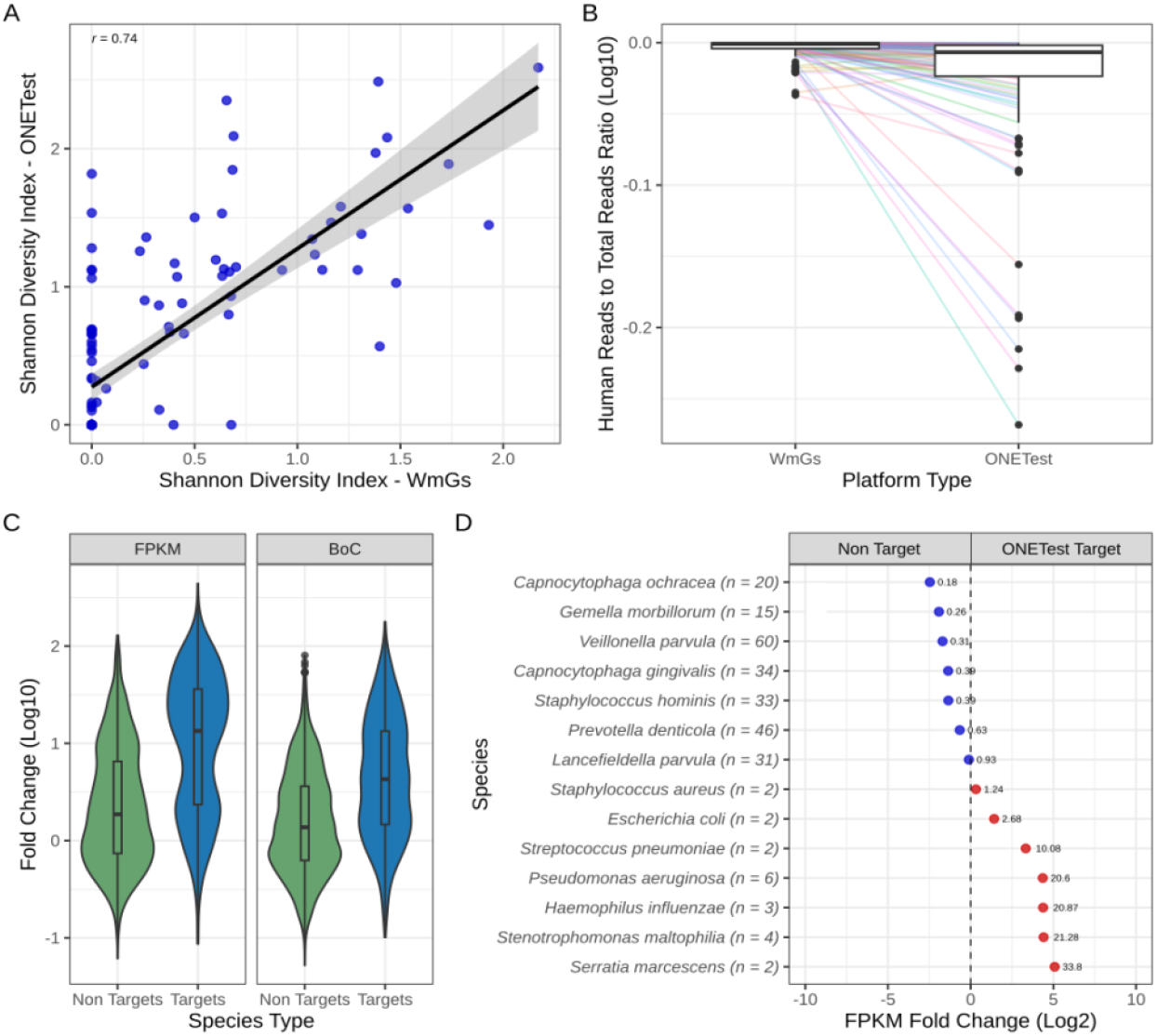
Comparative evaluation of ONETest Pathogenome and Whole-Metagenome Shotgun sequencing in BAL samples from cohort 1. **(A)** Scatter plot comparing Shannon diversity indices (capturing species richness and evenness) derived from OT (y-axis) and WmGS (x-axis) for 119 BAL samples in cohort 1. Data points represent individual samples, with a least-squares regression line and 95% confidence interval shaded in gray. Pearson correlation coefficient (*r*) is indicated. **(B)** Paired comparison of human reads/total clean reads ratio between OT and WmGS with dataset downsampled to match OT library sizes per sample (n = 119). Points represent individual samples connected by lines horizontal bars indicate medians with interquartile ranges. **(C)** Log10 fold-change in FPKM (left panel) and breadth of coverage (right panel) for targeted versus non-targeted species, comparing OT to library-size-matched WmGS data. Box plots display medians, interquartile ranges, and whiskers extending to 1.5 times the interquartile range. Individual points show per-sample species values. Species are categorized as non-targets (green) or targets (blue). **(D)** Log2 fold-change in FPKM for culture-positive species, comparing OT to full-depth WmGS data across the 119 BAL samples. Bars represent mean fold-changes per species, with error bars denoting standard deviations. n values indicate the number of culture-positive samples for each species. Red and blue bars denote OT-target and –non-target calls detected in this cohort, respectively. Species are listed in descending order of fold-change.

### Clinical performance (cohort 2)

#### OT distinguishes contaminants for high-specificity pathogens detection

To assess assay specificity and distinguish target signals from contaminants and background, 36 NTCs and nine HCC replicates were processed alongside 360 clinical BAL samples. These controls revealed detectable signals for 47 microbial species, among which 13 (Fig. S4A) are present in all HCC consistent with persistent background likely arising from reagents and/or sequencing. Other species displayed variable prevalence suggesting heterogeneous background sources across runs and sample processing. Median FPKM values varied widely by species – from 0.0002–0.09 in HCC, which informed subsequent filtering and empirical calling thresholds for clinical specimens (Fig. S4B).

#### OT detects and characterizes potential pathogens in 360-BAL samples (accuracy)

To benchmark OT against routine/AFB cultures, we analyzed cohort-2 specimens (Table S2) submitted for standard microbiology. Culture showed bacterial growth in 115/360 samples (32%), yielding 133 isolates (unique organisms identified in a given sample) across 23 species (Table 2). The most frequent bacterial pathogens (>5 isolates) were *P. aeruginosa* (n=37), *S. aureus* (n=19), *M. avium–intracellulare– chimaera* complex (MAC, n = 13), *S. maltophilia* (n=12), *H. influenzae* (n=7), *A. xylosoxidans* and *M. catarrhalis* (n=6 each). OT detected at least one culture-confirmed pathogen in 100/115 of positive cases (87%), with exact species/complex-level concordance in 89/115 (77%), comprising 11 polymicrobial and 78 single-pathogen cases. Among the remaining 11 culture-positive cases without exact concordance, culture reported ≥2 pathogens: in six cases, OT detected one or two of the cultured organism (FG073, FG090, FG160, FG187, FG267, FG334) but not the other; in three, the alternate OT detection was supported by MGDx or PCR when available (FG028, FG081, FG313); and in two, the OT-only signals could not be confirmed (Table S7, Fig. 2A). OT detected none of the cultured organisms in 15/115 culture positives where most OT false negatives occurred in specimens with sparse culture growth (Fig. S5).

**Table 2.** ONETest™ detection performance at species and sample levels in clinical-trial cohort-2 benchmarked against routine bacterial and AFB culture.

| Culture-confirmed bacterial spp. | Culture+ | TP | FN | FP | TN | accuracy | specificity | sensitivity | PPV | NPV | F1_score | Cohen's κ | Cohen's κ<br>(95% CI) |
| --- | --- | --- | --- | --- | --- | --- | --- | --- | --- | --- | --- | --- | --- |
| *Pseudomonas aeruginosa* | 37 | 34 | 3 | 23 | 300 | 0.93 | 0.93 | 0.92 | 0.60 | 0.99 | 0.72 | 0.68 | 0.57 – 0.80 |
| *Staphylococcus aureus* | 19 | 14 | 5 | 7 | 334 | 0.97 | 0.98 | 0.74 | 0.67 | 0.99 | 0.70 | 0.68 | 0.51 – 0.85 |
| *Mycobacterium avium* complex | 13 | 8 | 5 | 0 | 302 | 0.98 | 1.00 | 0.62 | 1.00 | 0.98 | 0.76 | 0.75 | 0.55 – 0.96 |
| *Stenotrophomonas maltophilia* | 12 | 11 | 1 | 8 | 340 | 0.98 | 0.98 | 0.92 | 0.58 | 1.00 | 0.71 | 0.70 | 0.51 – 0.88 |
| *Haemophilus influenzae* | 7 | 7 | 0 | 19 | 334 | 0.95 | 0.95 | 1.00 | 0.27 | 1.00 | 0.42 | 0.41 | 0.20 – 0.62 |
| *Achromobacter xylosoxidans* | 6 | 5 | 1 | 3 | 351 | 0.99 | 0.99 | 0.83 | 0.63 | 1.00 | 0.71 | 0.71 | 0.44 – 0.98 |
| *Moraxella catarrhalis* | 6 | 5 | 1 | 4 | 350 | 0.99 | 0.99 | 0.83 | 0.56 | 1.00 | 0.67 | 0.66 | 0.38 – 0.94 |
| *Escherichia coli* | 5 | 3 | 2 | 1 | 354 | 0.99 | 1.00 | 0.60 | 0.75 | 0.99 | 0.67 | 0.66 | 0.30 – 1.00 |
| *Klebsiella oxytoca* complex | 5 | 3 | 2 | 0 | 355 | 0.99 | 1.00 | 0.60 | 1.00 | 0.99 | 0.75 | 0.75 | 0.41 – 1.00 |
| *Klebsiella pneumoniae* | 5 | 5 | 0 | 2 | 353 | 0.99 | 0.99 | 1.00 | 0.71 | 1.00 | 0.83 | 0.83 | 0.60 – 1.00 |
| *Mycobacteroides abscessus* | 3 | 2 | 1 | 0 | 312 | 1.00 | 1.00 | 0.67 | 1.00 | 1.00 | 0.80 | 0.80 | 0.41 – 1.00 |
| *Pseudomonas fluorescens* complex | 2 | 1 | 1 | 0 | 358 | 1.00 | 1.00 | 0.50 | 1.00 | 1.00 | 0.67 | 0.67 | 0.05 – 1.00 |
| *Serratia marcescens* | 2 | 2 | 0 | 2 | 356 | 0.99 | 0.99 | 1.00 | 0.50 | 1.00 | 0.67 | 0.66 | 0.23 – 1.00 |
| *Streptococcus pneumoniae* | 2 | 2 | 0 | 9 | 349 | 0.98 | 0.97 | 1.00 | 0.18 | 1.00 | 0.31 | 0.30 | – 0.02 – 0.62 |
| *Acinetobacter baumannii* complex | 1 | 1 | 0 | 0 | 359 | 1.00 | 1.00 | 1.00 | 1.00 | 1.00 | 1.00 | 1.00 | 1.00 – 1.00 |
| *Bordetella parapertussis* | 1 | 1 | 0 | 2 | 357 | 0.99 | 0.99 | 1.00 | 0.33 | 1.00 | 0.50 | 0.50 | – 0.10 – 1.00 |
| *Burkholderia cepacia* | 1 | 1 | 0 | 0 | 359 | 1.00 | 1.00 | 1.00 | 1.00 | 1.00 | 1.00 | 1.00 | 1.00 – 1.00 |
| *Chryseobacterium indologenes* * | 1 | 0 | 1 | 0 | 359 | 1.00 | 1.00 | 0.00 | NA | 1.00 | 0.00 | 0.00 | 0.00 – 0.00 |
| *Enterobacter cloacae* complex | 1 | 1 | 0 | 0 | 359 | 1.00 | 1.00 | 1.00 | 1.00 | 1.00 | 1.00 | 1.00 | 1.00 – 1.00 |
| *Mycobacterium gordonae* | 1 | 0 | 1 | 0 | 314 | 1.00 | 1.00 | 0.00 | NA | 1.00 | 0.00 | 0.00 | 0.00 – 0.00 |
| *Mycobacterium kansasii* | 1 | 0 | 1 | 0 | 314 | 1.00 | 1.00 | 0.00 | NA | 1.00 | 0.00 | 0.00 | 0.00 – 0.00 |
| *Proteus mirabilis* | 1 | 1 | 0 | 0 | 359 | 1.00 | 1.00 | 1.00 | 1.00 | 1.00 | 1.00 | 1.00 | 1.00 – 1.00 |
| *Streptococcus pyogenes* | 1 | 0 | 1 | 0 | 359 | 1.00 | 1.00 | 0.00 | NA | 1.00 | 0.00 | 0.00 | 0.00 – 0.00 |
| **Micro-averaged total isolates** | **133** | **107** | **26** | **80** | **7528** | 0.99 | 0.99 | 0.80 | 0.57 | 1.00 | 0.67 | 0.66 | 0.60 – 0.73 |
| **Total samples** | **115** | **95** | **20** | **76** | **169** | 0.73 | 0.69 | 0.83 | 0.56 | 0.90 | 0.66 | 0.46 | 0.38 – 0.56 |

\* Species not included in the ONETest Pathogenome panel. *Mycobacterium avium* complex (MAC) represents *Mycobacterium avium*–*intracellulare*–*chimaera* species. NA, not applicable (undefined when TP+FP=0). Cohen's κ was interpreted per Landis and Koch (1977). Micro-averaged performance metrics were calculated by pooling TP, FP, FN, and TN across all species-level isolates/calls (i.e., aggregated confusion matrices across species rather than sample-level performance). *Mycobacterium* spp. performance metrics were calculated using the subset of samples with available AFB culture results; all other species were evaluated across the full cohort.

**Figure 2.**
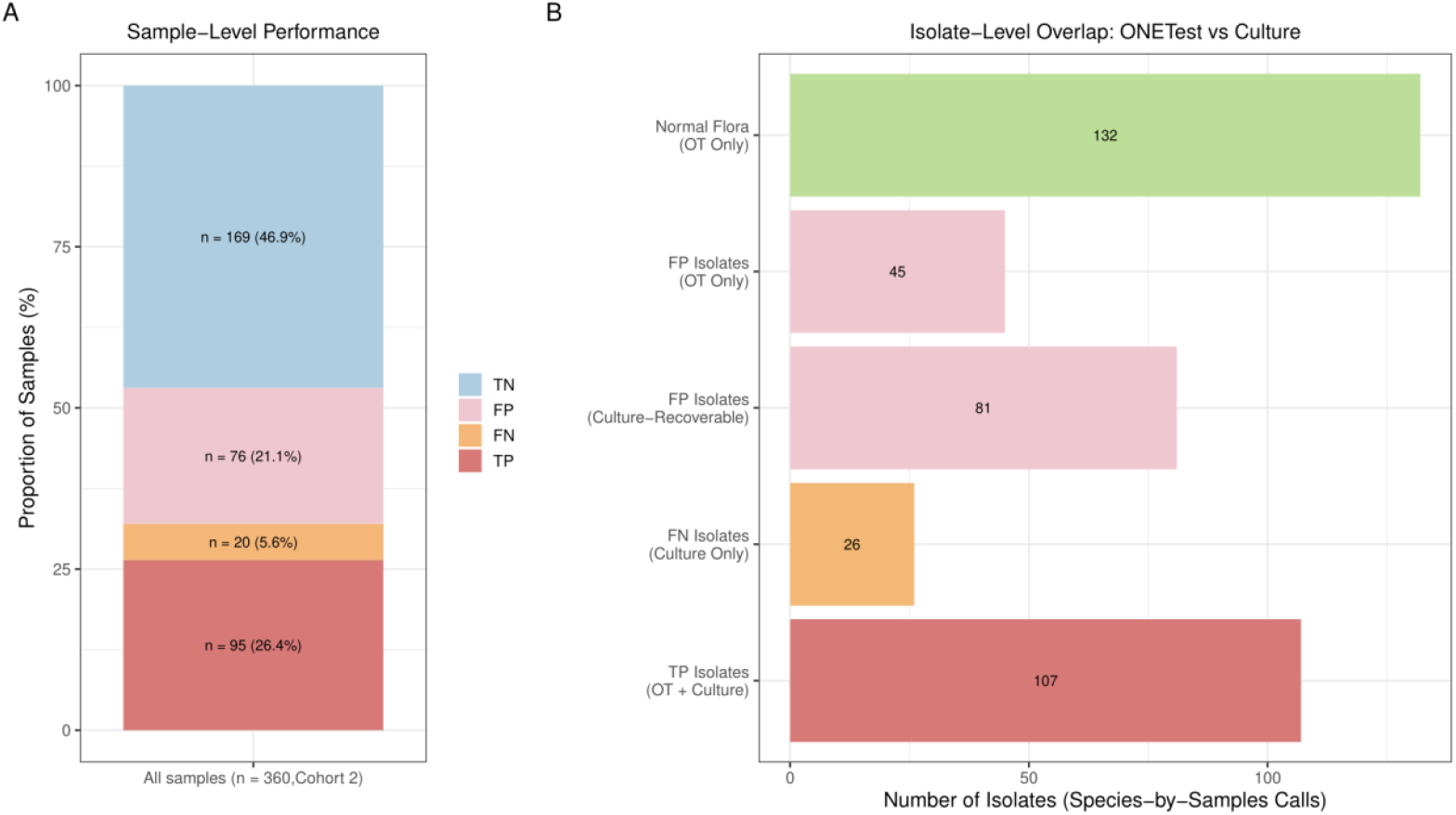
ONETest™ detection performance versus culture at specimen- and isolate-level in the clinical trial cohort 2. **(A) Sample-level detection performance for 360 BAL specimens**. Each sample classified as true positive (TP: OT detected ≥1 culture-positive pathogen 95/360), false negative (FN: culture grew ≥1 pathogen not detected by OT, 20/360), false positive (FP: OT positive with no culture-proven pathogen, 76/360), or true negative (TN: neither method detected pathogens beyond normal flora, 169/360). When multiple outcomes were possible for a specimen, TP and FN status were prioritized over FN and FP, respectively. Bars show the proportion of samples/cases in each category. **(B) Isolate-level overlap between OT and culture across all species-sample combinations (isolates/calls) in cohort 2**. Bars show isolates detected by both methods (TP isolates), by culture only (FN isolates), by OT additional corresponding to species cultured elsewhere in the cohort (FP isolates – cultured-recoverable species), and for species not recovered by culture in this cohort (FP isolates – OT only), and OT calls survived and reported as normal flora after the clinical-detection thresholding. Bar lengths and labels indicate isolate counts.

For most prevalent culture-detected pathogens, OT showed high accuracy and specificity for species detection (Table 2). Among bacteria with >5 culture-positive isolates (excluding MAC), accuracy ranged from 93% to 99% and specificity from 93% to 100%. MAC showed a sensitivity of 62% and an accuracy of 98%. Following NTM–OT discordance, 16S-rDNA qPCR was performed on a subset of concordant and discordant specimens. Both OT negatives/culture positives (FG-267 and FG-250) were negative by qPCR, whereas the OT positive/culture positive FG-258 showed consistent qPCR amplification (Table S8). Micro-averaged across all isolates/species, OT achieved an overall sensitivity of 0.80, specificity of 0.99, and overall agreement (Cohen’s κ) of 0.66. For prevalent species (n≥5), F1-scores (0.42–0.83) and κ values (0.41–0.83) indicated moderate-to-substantial agreement, with common pathogens such as *P. aeruginosa* and *S. maltophilia* showing high sensitivity (92% each) and strong F1-scores. Although the F1-scores for *H. influenzae* (0.42, likely influenced by false positives) and *M. catarrhalis* (0.67) were lower than common species, OT still achieved high sensitivity (100% and 83%, respectively) and maintained moderate-to-substantial agreement. Rarer species exhibited more variable performance, likely driven by low prevalence (Table 2). At the sample level, OT achieved an overall accuracy of 73% and sensitivity of 83% for culture positives and specificity of 69% for culture negatives (Table 2, Fig. 2A).

#### OT detects additional species beyond culture in 360-BAL samples

OT identified 258 additional isolates not recovered by routine bacterial/AFB culture in cohort 1 (Table S7, Fig. 2B). These isolates were classified as either LRT-associated pathogens or taxa consistent with normal flora/upper-airway commensals (Fig. S6). Of 258 additional isolates, 126 (49%) were LRT-associated organisms, including 81 culture-recoverable isolates (species cultured elsewhere in the cohort) and 45 OT-only isolates. These LRT-associated OT calls occurred in 104/360 specimens, comprising 76 OT-positive/culture-negative specimens and 28 culture positives with additional OT co-detections. Among the 76 OT-positive/culture-negative specimens, 94 isolates were reported (additive yield, 21% [76/360]), corresponding to a mean of 1.24 reportable organisms per OT-positive/culture-negative specimen; these comprised 62 culture-recoverable and 32 OT-only detections (Fig. S6). Remaining 132 isolates (51.4%) comprised 26 taxa consistent with normal flora/upper-airway commensals; culture results reported as “normal flora” were treated as culture-negative, and isolates classified as normal under OT reporting schemes were treated as OT-negative for performance analyses.

We orthogonally assessed OT-additional isolates in a subset BAL specimen (n=92) also tested by the BioFire Pneumonia (PN) panel (Figs. S2, S7). Of 70/81 culture-recoverable additional isolates tested by PN, 58 (82.9%) were PN evaluable. Among evaluable isolates, 28/58 (48.3%) were PN-positive for the same species. Most OT– PN discordant isolates involved *P. aeruginosa*; excluding *P. aeruginosa*, 72% of remaining isolates were PN-positive (Fig. S7), with high concordance for *H. influenzae* (13/16) and *S. aureus* (4/5). Among 45 OT-only isolates, only two (*Streptococcus agalactiae*) were PN-covered, and both were PN-positive. Overall, 60 OT additional isolates across 54 specimens were PN-evaluable, yielding 30 PN-concordant detections in 27 specimens: 17/92 (18.5%) were culture-negative for the corresponding organism and 10 represented PN-supported co-detections in culture-positive specimens. Extrapolated to the full cohort, 27/360 (7.5%) BAL specimens gained ≥1 additional PN-supported microbiologic findings.

#### OT versus MGDx for culture-confirmed pathogen detection

We benchmarked OT against a sequencing-based comparator, analysing a subset of cohort-2 (n=66) that was also evaluated by MicroGenDx(MGDx) (Fig. 3, Table S9). Routine culture was positive in 22/66 samples, yielding 23 culture-confirmed isolates, including one polymicrobial specimen. Overall, 47/66 (71%) samples were positive for at least one organism by at least one method, yielding 82 organism detections across the three assays. Across all detections, MGDx identified 58/82 (70%) isolates spanning 21 species, OT identified 41/82 (50%) isolates spanning 17 species, and culture identified 22/82 (27%) isolates spanning 10 species. Using culture as the reference standard, 10/22 (45%) isolates were detected by all three methods, 8/22 (35%) were detected by OT but not MGDx, 2/22 (9%) were detected by MGDx but not OT, and 2/22 (9%) were missed by both sequencing assays. Accordingly, OT detected 19/22 (86% [95% CI:67–95%]) culture-confirmed isolates, compared with 12/22 (54% [95% CI: 34–75%]) detected by MGDx, indicating higher sensitivity for recovery of culture-confirmed pathogens in this BAL subset. In addition, OT and MGDx shared nine concordant detections not recovered by culture and two concordant false negatives (culture-positives missed by both sequencing assays), supporting overlap in some additional culture negatives and shared misses.

**Figure 3.**
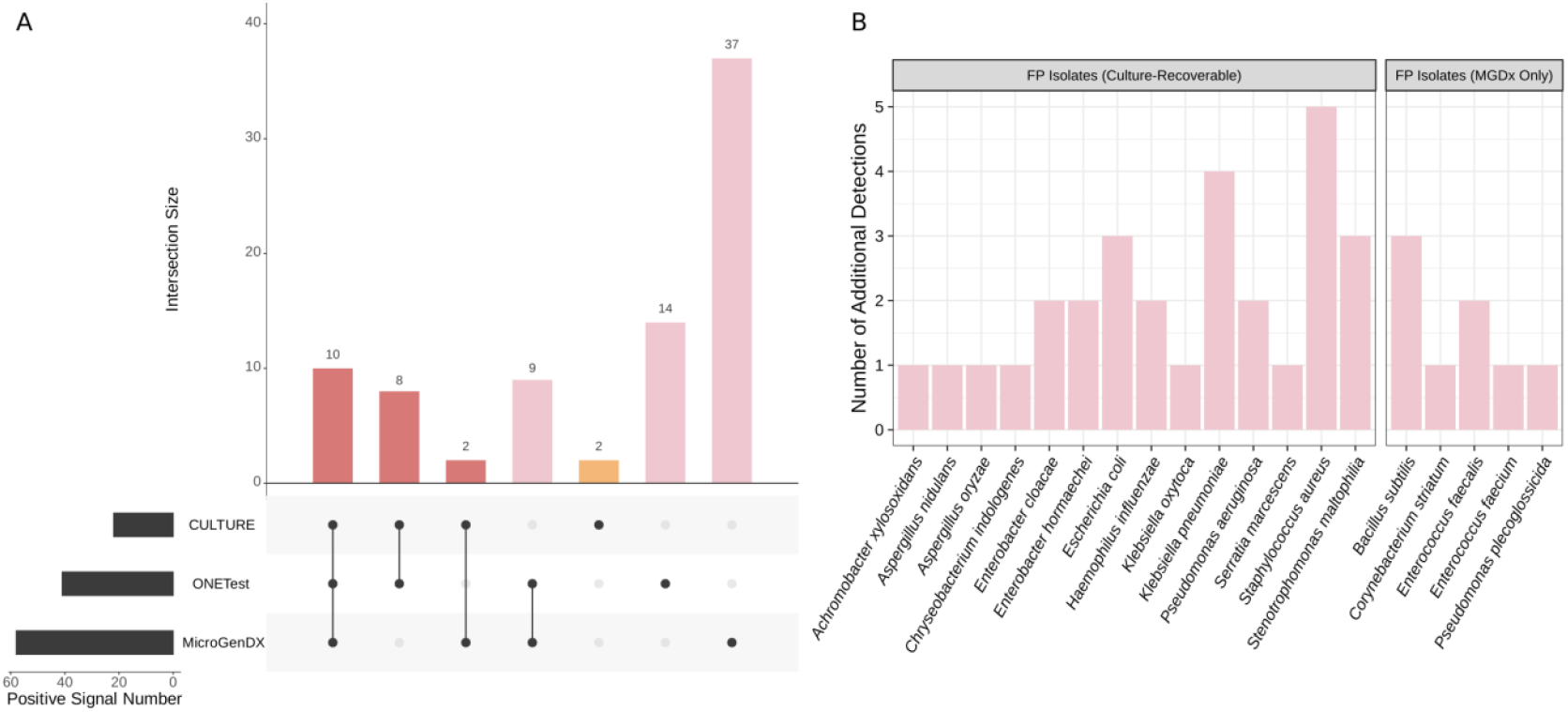
Comparative detection performance of ONETest™ (Targeted WmGS) and MicroGenDx (Amplicon-based NGS) against culture in BAL Samples. **(A)** Intersection of positive detections across culture, OT, and MGDx in a subset of 66 BAL samples from cohort 2 stratified by the number of positive detections per method. Bars represent the total number of positive signals for each method, highlighting OT’s higher detection rate compared to MGDx while maintaining closer alignment with culture positives. **(B)** Distribution of additional species detected beyond culture by MGDx, categorized as culture-recoverable species (left) and additional species only detected by MGDx (right), with bar heights representing the number of detections per species.

### LDT performance/validation (cohort 3)

To evaluate OT as an LDT, we analyzed 43 BAL-cohort 3 specimens (Table 3). Culture and AFB testing identified 40/43 positive cases, yielding 55 bacterial and fungal isolates, including polymicrobial specimens (Table 3, Fig. 4A). The most frequently recovered organisms were the MAC complex (n = 10), *P. aeruginosa* (n = 9), *A. xylosoxidans* and *B. cepacia* complex (each n = 4). Remaining isolates comprised less frequent bacterial and fungal spp. (≤3 isolates each,). At species level, OT detected 41/55 culture-confirmed isolates (Fig. 4B), corresponding to a micro-averaged sensitivity of 0.75, specificity of 0.98, and agreement of 0.73 Cohen’s κ with culture. The MAC complex accounted for the largest number of culture-positive isolates in this cohort and remained challenging, with a sensitivity of 60% (6/10) but high specificity (97%) and an overall accuracy of 88% (Cohen’s κ = 0.63; 95% CI, 0.35–0.92). For *P. aeruginosa*, OT achieved 89% (8/9) sensitivity, 85% specificity. Highly concordant results were observed for several less frequent but clinically relevant bacterial spp., including *A. xylosoxidans* and *B. cepacia* complex, for which OT achieved high to perfect sensitivity and specificity (≥75– 100%) and substantial agreement (κ ≈ 0.77–0.84). In contrast, fungal pathogens exhibited greater variability. *A. fumigatus* and *Coccidioides posadasii* were detected in 2/3 culture-positive isolates (66% sensitivity), whereas other culture-confirmed fungal species (n=4) were not detected by OT in this small sample set, yielding nominal sensitivities of 0–50% despite uniformly high specificity (1.00). For very low-prevalence taxa (n ≤ 2), point estimates were frequently 100% but confidence intervals were wide due to limited sample size.

**Table 3.** ONETest™ detection performance at species and sample levels in validation cohort-3 benchmarked against routine bacterial and AFB culture.

| Bacterial and fungal species | Culture + | TP | FN | FP | TN | accuracy | specificity | sensitivity | PPV | NPV | F1_score | Cohen's κ | Cohen's κ (95% CI) |
| --- | --- | --- | --- | --- | --- | --- | --- | --- | --- | --- | --- | --- | --- |
| *Mycobacterium avium complex* | **10** | 6 | 4 | 1 | **31** | 0.88 | 0.97 | 0.60 | 0.86 | 0.89 | 0.71 | 0.63 | 0.35 – 0.92 |
| *Pseudomonas aeruginosa* | 9 | 8 | 1 | 5 | 29 | 0.86 | 0.85 | 0.89 | 0.62 | 0.97 | 0.73 | 0.64 | 0.38 – 0.90 |
| *Achromobacter xylosoxidans* | 4 | 4 | 0 | 2 | 37 | 0.95 | 0.95 | 1.00 | 0.67 | 1.00 | 0.80 | 0.77 | 0.48 – 1.00 |
| *Burkholderia cepacia complex* | 4 | 3 | 1 | 0 | 39 | 0.98 | 1.00 | 0.75 | 1.00 | 0.98 | 0.86 | 0.84 | 0.55 – 1.00 |
| *Chryseobacterium indologenes* * | 3 | 2 | 1 | 0 | 40 | 0.98 | 1.00 | 0.67 | 1.00 | 0.98 | 0.80 | 0.79 | 0.39 – 1.00 |
| ***Aspergillus fumigatus*** | 2 | 1 | 1 | 0 | 41 | 0.98 | 1.00 | 0.50 | 1.00 | 0.98 | 0.67 | 0.66 | 0.03 – 1.00 |
| *Enterobacter cloacae complex* | 2 | 2 | 0 | 0 | 41 | 1.00 | 1.00 | 1.00 | 1.00 | 1.00 | 1.00 | 1.00 | 1.00 – 1.00 |
| *Klebsiella oxytoca complex* | 2 | 2 | 0 | 0 | 41 | 1.00 | 1.00 | 1.00 | 1.00 | 1.00 | 1.00 | 1.00 | 1.00 – 1.00 |
| *Moraxella catarrhalis* | 2 | 2 | 0 | 0 | 41 | 1.00 | 1.00 | 1.00 | 1.00 | 1.00 | 1.00 | 1.00 | 1.00 – 1.00 |
| *Mycobacteroides abscessus* | **2** | 2 | 0 | 0 | 40 | 1.00 | 1.00 | 1.00 | 1.00 | 1.00 | 1.00 | 1.00 | 1.00 – 1.00 |
| ***Paecilomyces sp.*** | 2 | 0 | 2 | 0 | 41 | 0.95 | 1.00 | 0.00 | NA | 0.95 | 0.00 | 0.00 | 0.00 – 0.00 |
| *Staphylococcus aureus* | 2 | 2 | 0 | 2 | 39 | 0.95 | 0.95 | 1.00 | 0.50 | 1.00 | 0.67 | 0.64 | 0.20 – 1.00 |
| *Stenotrophomonas maltophilia* | 2 | 2 | 0 | 2 | 39 | 0.95 | 0.95 | 1.00 | 0.50 | 1.00 | 0.67 | 0.64 | 0.2 – 1.00 |
| ***Aspergillus niger*** | 1 | 0 | 1 | 0 | 42 | 0.98 | 1.00 | 0.00 | NA | 0.98 | 0.00 | 0.00 | 0.00 – 0.00 |
| ***Aspergillus spp.*** | 1 | 0 | 1 | 0 | 42 | 0.98 | 1.00 | 0.00 | NA | 0.98 | 0.00 | 0.00 | 0.00 – 0.00 |
| ***Coccidioides posadasii*** | 1 | 1 | 0 | 0 | 42 | 1.00 | 1.00 | 1.00 | 1.00 | 1.00 | 1.00 | 1.00 | 1.00 – 1.00 |
| *Haemophilus influenzae* | 1 | 1 | 0 | 3 | 39 | 0.93 | 0.93 | 1.00 | 0.25 | 1.00 | 0.40 | 0.38 | –0.16 – 0.91 |
| *Mycobacterium gordonae* | **1** | 0 | 1 | 0 | 41 | 0.98 | 1.00 | 0.00 | NA | 0.98 | 0.00 | 0.00 | 0.00 – 0.00 |
| *Mycobacterium tuberculosis* | **1** | 1 | 0 | 0 | 41 | 1.00 | 1.00 | 1.00 | 1.00 | 1.00 | 1.00 | 1.00 | 1.00 – 1.00 |
| *Pasteurella multocida* | 1 | 1 | 0 | 0 | 42 | 1.00 | 1.00 | 1.00 | 1.00 | 1.00 | 1.00 | 1.00 | 1.00 – 1.00 |
| *Pseudomonas stutzeri* * | 1 | 1 | 0 | 0 | 42 | 1.00 | 1.00 | 1.00 | 1.00 | 1.00 | 1.00 | 1.00 | 1.00 – 1.00 |
| *Streptococcus pneumoniae* | 1 | 0 | 1 | 0 | 42 | 0.98 | 1.00 | 0.00 | NA | 0.98 | 0.00 | 0.00 | 0.00 – 0.00 |
| **Micro-averaged total isolates** | **55** | **41** | **14** | **15** | **873** | 0.97 | 0.98 | 0.75 | 0.73 | 0.98 | 0.74 | 0.72 | 0.61 – 0.81 |
| **Total samples** | **40** | **34** | **6** | **1** | **2** | 0.84 | 0.67 | 0.85 | 0.97 | 0.25 | 0.91 | 0.29 | –0.08 – 0.66 |

\* Species not included in the ONETest Pathogenome panel. *Mycobacterium avium complex* (MAC) represents *Mycobacterium avium–intracellulare–chimaera* species. NA, not applicable (undefined when TP+FP=0). Cohen's κ was interpreted per Landis and Koch (1977). Micro-averaged performance metrics were calculated by pooling TP, FP, FN, and TN across all species-level isolates/calls (i.e., aggregated confusion matrices across species rather than sample-level performance). *Mycobacterium spp.* performance metrics were calculated using the subset of samples with available AFB culture results; all other species were evaluated across the full cohort. Fungal species are bolded.

**Figure 4.**
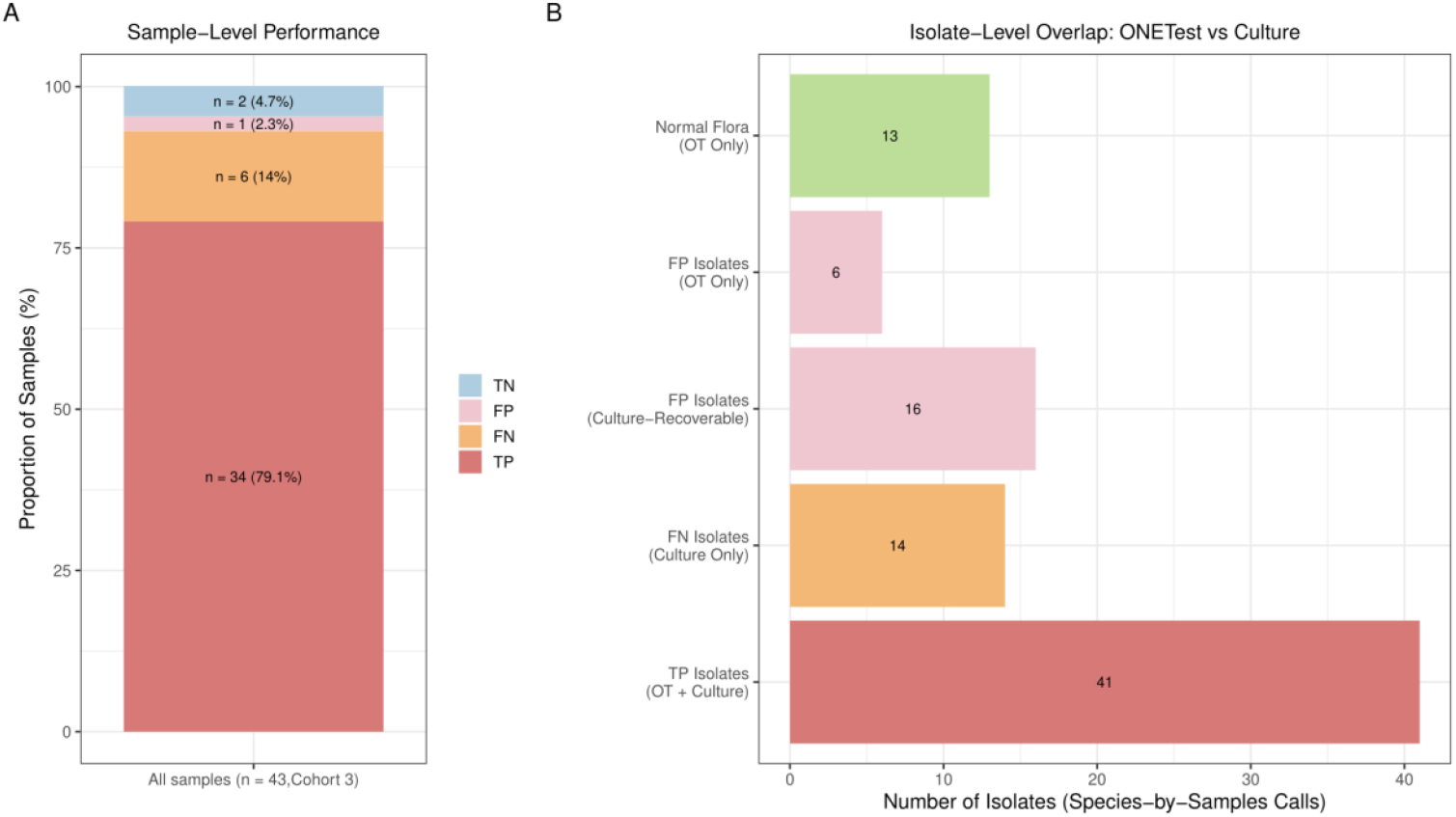
ONETest™ detection performance versus culture at specimen- and isolate-level in the LDT validation cohort 3. **(A) Sample-level detection performance of 43 BAL specimens**. Each specimen was categorized as true positive (TP: OT detected ≥1 culture-positive pathogen, 34/43), false negative (FN: culture grew ≥1 pathogen not detected by OT, 6/43), false positive (FP: OT positive with no culture-proven pathogen,1/43), or true negative (TN: neither method detected pathogens beyond normal flora, 2/43). When multiple outcomes were possible for a specimen, TP status was prioritized over FN. Bar height represents the proportion of specimens in each category. (**B) Isolate-level overlap between ONETest and culture across all species-sample combinations (isolates) in cohort 3**. Bars show isolates detected by both methods (TP isolates, n = 41), culture-positive isolates missed by OT (FN isolates, n = 14), OT-only detections corresponding to species cultured in other specimens from the same cohort (FP isolates – cultured species, n = 16), OT-only detections corresponding to additional target species not recovered by culture in this cohort (FP isolates – additional species, n = 6), and ONETest detections reported as normal flora in BAL specimens (normal flora, n = 13). Bar lengths correspond to isolate counts, and numeric labels indicate the absolute number of isolates in each group.

At the sample level, OT identified at least one culture-confirmed pathogen in 34/40 culture-positive specimens (85%), with one OT-positive/culture-negative specimen and two specimens negative by both methods (Table 3, Fig. 4A). This corresponded to an overall accuracy of 84%, sensitivity of 85% for culture-positive samples, and specificity of 67% for culture-negative samples. Given only three culture-negative specimens, specificity and NPV estimates were imprecise.

OT detected 16 culture-recoverable additional isolates across 15/43 specimens. Of these, 9/16 (56.3%) were covered by the PN panel. Among PN-evaluable isolates, 8/9 were PN-positive and 1/9 was negative (Fig. S7). In addition, among six OT-only species, two (adenovirus and *E. coli*) were confirmed by both OT and the PN panel. In total, 11 OT-only additional isolates (9 culture-recoverable and 2 OT-only) were orthogonally evaluated across ten OT-only positive specimens, with 10/11 calls concordant between OT and PCR-based testing. Among these ten specimens, one was culture-negative for any pathogen, while nine were culture-positive for a different organism, consistent with polymicrobial detection not resolved by routine culture.

## DISCUSSION

Despite advances in culture workflows and syndromic PCR, BAL-based evaluation of suspected pneumonia often yields results that are delayed, negative, or difficult to interpret, particularly after antibiotics and in immunocompromised hosts^8^. The key challenge is not simply detecting microbial nucleic acid, but delivering timely, contamination-aware results that help distinguish infection from colonization in a host-rich, non-sterile specimen. NGS-based approaches (WmGS, metatranscriptomics, and TmGS) have therefore emerged as potential adjuncts^9^, but adoption has been limited by host-background dilution, workflow variability, and analytic complexity along with the need for control-informed interpretation so that additional detections do not translate into unnecessary antimicrobial exposure^5,8,9^.

Against this backdrop, we evaluated OT across analytical, technical, and clinical BAL cohorts to define performance characteristics relevant to routine respiratory diagnostics. OT achieved a clinically relevant reporting threshold in patient-matrix BAL and, relative to WmGS, preserved overall community structure while improving sequencing efficiency through on-target enrichment. In a 360-sample retrospective BAL cohort, OT showed high concordance with routine bacterial culture and good agreement with AFB culture at both species and sample levels, and it identified additional organisms not recovered by culture. In a tested subset, OT also showed higher sensitivity than an amplicon-based sequencing comparator for recovery of culture-confirmed targets. OT met performance expectations in a CAP-accredited LDT validation. Together, these data support OT as an adjunct to conventional microbiology for BAL evaluation when results are interpreted in clinical context^5,10^.

Clinically, the question is whether an NGS adjunct can change bedside decisions. The performance profile observed here suggests OT may be most useful as a complement to routine microbiology in patients with high pretest probability of infection but limited yield from standard testing (e.g., severe pneumonia in the ICU, immunocompromised hosts, structural lung disease with prior antibiotics, suspected polymicrobial infection, or concern for slow-growing/fastidious organisms). Concordant OT detections may support earlier tailoring of antimicrobial therapy when aligned with the clinical picture, while OT-negative results for common bacterial pathogens may provide additional support for de-escalation when paired with clinical trajectory and imaging. Conversely, OT-positive/culture-negative results should be framed as additional detections rather than definitive infection and interpreted alongside semiquantitative signal, commensal annotation, antibiotic exposure, and orthogonal evidence (e.g., multiplex PCR) to minimize unnecessary antimicrobial treatment.

The LoD range reported here (2 ×10^3^–10^4^ CFU/mL) reflects a context-aware reporting threshold in BAL rather than an absolute analytical limit. This approach accounts for background signal and emphasizes biological relevance in a non-sterile specimen. Notably, the LoD range overlaps quantitative BAL culture thresholds (typically ≥10^4^ CFU/mL) commonly used to support infection rather than colonization or upper-airway contamination^11^, consistent with the analytical sensitivity framework described by Chiu and Miller (2019). Similar BAL LoD ranges have been reported in other respiratory NGS evaluations^10,12,13^. In contrast, lower LoDs (e.g., 50–450 CFU/mL)^14^ often reflect synthetic matrices and different reporting rules, limiting cross-platform clinical comparability.

OT uses hybrid capture to approximate WmGS community profiles while reducing sequencing burden. In cohort 1, OT and WmGS showed concordant community structure and species richness, and at comparable depth OT reduced host background while increasing reads assigned to culture-confirmed pathogens. As reported for other capture-based approaches, enrichment varies by organism class and genome features: virus-focused designs can yield large gains in viral reads at reduced depth, whereas multi-kingdom panels often show more modest bacterial and fungal gains in clinical specimens^15–17^. Consistent with a multi-kingdom design, OT increased per-pathogen signal (up to 33-fold) relative to full-depth WmGS for most culture-confirmed pathogens (Fig. 1D), including select slow-growing/fastidious organisms (e.g., culture-positive *M. kansasii* and *M. abscessus*) that were missed by WmGS or amplicon TmGS (Tables S7, S10). These gains at lower sequencing depth support improved feasibility for implementation.

In the 360-sample retrospective BAL cohort, OT showed high concordance with routine culture for common pathogens and strong sample-level rule-out performance (NPV 90%). When culture is the primary comparator in non-sterile BAL, lower apparent PPV and specificity are expected because antibiotic exposure and low organism burden can reduce culture yield and sequencing can detect low-level signal consistent with colonization or antibiotic-suppressed infection not recovered by culture. OT detected at least one culture-confirmed target in 88% of culture-positive BALs, with sample-level sensitivity of 83%, specificity of 69%, and NPV of 90% (κ = 0.47). At the isolate level, OT maintained very high micro-averaged specificity (0.99) with sensitivity of 0.81 (F1 = 0.67; κ = 0.67). Accordingly, comparisons across studies are most interpretable when they use the same unit of analysis (sample vs organism-call) and the same reference standard (culture-only vs composite adjudication)^5,8^.

Discordance was most pronounced for NTM and should be interpreted in the context of specimen workflow. NTM testing was performed on residual BAL after routine hematology, where available volume was limited and pre-extraction concentration was not feasible. PCR testing of OT-negative/culture-positive specimens did not detect mycobacterial DNA, suggesting that discordance in this cohort reflected volume-constrained DNA recovery – a major determinant of sensitivity^18^, rather than enrichment or sequencing failure. This interpretation is supported by internal mycobacteria-spiking experiments in BAL matrices, which showed culture-concordant OT performance when sufficient material was available (Fig. S8). For laboratories implementing sequencing-based diagnostics, these results underscore the importance of ensuring adequate specimen volume (and, when available, concentration strategies) for NTM recovery rather than relying on residual aliquots. Similarly, review of cohort-1 discordances for *S. aureus* found no organism-attributable reads in either WmGS or OT in 4/5 culture-positive/OT-negative specimens; the corresponding cultures were reported as “few *S. aureus*,” consistent with low organism burden rather than thresholding effects.

Beyond recapitulating culture-positive findings, OT provided incremental microbiologic information after commensal filtering, with an additive 20.8% specimen-level yield of OT-only target detections. Additional detections of this magnitude are common in respiratory NGS studies^5,12,19^, and likely reflect, in part, organisms not recovered by culture (including DNA viruses). Orthogonal multiplex PCR and available clinical documentation supported the plausibility of a subset of OT-only detections, but discordance particularly for organisms such as *P. aeruginosa* highlights a core challenge in non-sterile BAL: increased analytical sensitivity can surface low-level detections consistent with colonization, aspiration, or background signal rather than causative infection. Accordingly, OT-only detections are best interpreted in context and, where feasible, alongside contamination-aware thresholds and complementary diagnostic evidence^5,8,20^.

Comparison with an amplicon-based TmGS assay further clarifies OT’s positioning in BAL workflows. In the MGDx-evaluated subset, both sequencing assays reported organisms beyond routine culture, but OT recovered a higher proportion of culture-confirmed pathogens than MGDx (82.6% vs 52.2%). This pattern is consistent with differences in enrichment chemistry: amplicon approaches depend on primer binding at predefined loci and can be vulnerable to sequence variation, whereas hybrid capture distributes baits across many loci and can tolerate bait–target divergence, supporting detection across divergent lineages^16,21^. OT also detected organisms beyond the enumerated target list (e.g., *C. indologenes* and *P. stutzeri* in Table 3), underscoring that nominal “panel size” is an imperfect proxy for functional coverage, which depends on enrichment strategy and genomic breadth per organism^22^. Overall, these findings align with prior studies showing hybrid capture can improve recovery of culture-confirmed pathogens in BAL while supporting species-level confirmation and characterization^14,22^.

In cohort 3 (CAP-accredited LDT validation), OT showed reproducible performance under a standardized clinical laboratory workflow, supporting transportability of empirically defined detection rules. As with other BAL sequencing approaches^5,8,12^, results remain most interpretable when integrated with clinical context, particularly for low-burden or low-virulence organisms. For example, *M. gordonae*, which contributed to discordant calls, is often encountered in settings consistent with colonization or water-associated contamination; therefore, a culture-positive/OT-negative discordance may reflect non-disease isolation rather than a true false negative when there is no corroborating evidence or repeat positive cultures. OT identified ≥1 culture-confirmed organism in 34/40 culture-positive specimens (sensitivity 0.85; Table 3) with substantial isolate-level agreement (κ≈0.72). Apparent PPV was higher than in cohort 2, but interpretation is limited because cohort 3 was culture-enriched and included only three culture-negative specimens. OT showed strongest concordance for common cultured bacteria (e.g., *P. aeruginosa, A. xylosoxidans, B. cepacia* complex), while NTM remained biology- and burden-limited (e.g., MAC sensitivity 60%). For fungi, small denominators preclude stable estimates and motivate larger adjudicated cohorts. Continued accumulation of prospectively collected BAL specimens should enable refinement of species-specific baselines and thresholds, particularly for low-burden organisms such as NTM and molds under a controlled update process. This aligns with the broader NGS literature in which fungal detection is heterogeneous and specimen-dependent: plasma microbial cfDNA shows high specificity but modest sensitivity for invasive aspergillosis (∼38–44%), whereas BAL mNGS studies of invasive pulmonary aspergillosis report higher sensitivities (∼80–93% at study-specific thresholds)^23,24^.

Several limitations warrant consideration. First, analytical sensitivity was evaluated in a limited set of organisms at a fixed sequencing depth; performance may differ for other taxa and alternative reporting rules. Second, the clinical evaluation used residual, stored BAL aliquots processed after routine hematology. Limited volume and the inability to concentrate specimens likely reduced sensitivity for both OT and other sequencing approaches compared with standard microbiology workflows. Third, because routine culture served as the primary reference, performance metrics reflect agreement with culture rather than a composite clinical truth, and OT-positive/culture-negative detections cannot be adjudicated without systematic orthogonal testing and clinical correlation. Finally, estimates for less prevalent taxa, particularly fungi, were based on small denominators and require larger cohorts with adjudicated endpoints.

In summary, this multi-cohort evaluation shows that ONETest™ PathoGenome can provide broad pathogen detection in BAL with high concordance to routine bacterial and AFB culture and an additive yield that includes orthogonally supported detections not recovered by culture or an amplicon-based sequencing comparator. When paired with contamination-aware interpretation, these data support OT as a complementary diagnostic approach for complex, polymicrobial, or culture-limited presentations, including patients at increased risk of opportunistic infections. Prospective studies with adjudicated endpoints are needed to quantify incremental clinical impact and refine thresholds that better distinguish causation from colonization in lower-respiratory specimens.

## Author Approvals

All authors have reviewed and approved the final version of this manuscript. This work is a preprint and has not been published elsewhere.

## Author Contribution

**SMA:** Supervision, Conceptualization, Data Curation, Formal Analysis, Methodology, Supervision, Visualization, Writing - Original Draft; Writing - Review and Editing

**HDN:** Data Curation, Methodology, Software, Formal Analysis, Visualization, Writing - Review and Editing

**HD:** Data Curation, Methodology, Formal Analysis, Visualization

**NM:** Investigation

**BSK**.: Conceptualization, Investigation, Methodology, Project Administration, Supervision, Validation

**HJH:** Investigation, Validation

**GS:** Data Curation, Formal analysis, Funding Acquisition, Supervision

**TP:** Conceptualization, Writing - Review and Editing

**CK:** Conceptualization, Writing - Review and Editing

**PS:** Methodology, Project Administration, Resources, Supervision

**MAQ:** Conceptualization, Funding Acquisition, Project Administration, Resources, Supervision, Writing – review & editing

**KHR:** Supervision, Conceptualization, Writing - Review and Editing, Data Curation

## Acknowledgement

We thank Alex Adrian for assistance with the data analysis pipeline.

## References

1. Li, M., Liu, M. & Liu, J. Trends in the Mortality, Deaths, and Aetiologies of Lower Respiratory Infections Among 204 Countries from 1991 to 2021: An Updated Systematic Study. Viruses 17, 892 (2025).

2. Jain, S. et al. Community-Acquired Pneumonia Requiring Hospitalization among U.S. Adults. New England Journal of Medicine 373, 415–427 (2015).

3. Zhang, H. et al. Detection of Viruses by Multiplex Real-Time Polymerase Chain Reaction in Bronchoalveolar Lavage Fluid of Patients with Nonresponding Community-Acquired Pneumonia. Canadian Respiratory Journal 2020, 8715756 (2020).

4. Blauwkamp, T. A. et al. Analytical and clinical validation of a microbial cell-free DNA sequencing test for infectious disease. Nat Microbiol 4, 663–674 (2019).

5. Chiu, C. Y. & Miller, S. A. Clinical metagenomics. Nat Rev Genet 20, 341–355 (2019).

6. Costello, M. et al. Characterization and remediation of sample index swaps by non-redundant dual indexing on massively parallel sequencing platforms. BMC Genomics 19, 332 (2018).

7. Janda, J. M. & Abbott, S. L. 16S rRNA Gene Sequencing for Bacterial Identification in the Diagnostic Laboratory: Pluses, Perils, and Pitfalls. J Clin Microbiol 45, 2761–2764 (2007).

8. Simner, P. J., Miller, S. & Carroll, K. C. Understanding the Promises and Hurdles of Metagenomic Next-Generation Sequencing as a Diagnostic Tool for Infectious Diseases. Clin Infect Dis 66, 778–788 (2018).

9. Schlaberg, R. et al. Validation of Metagenomic Next-Generation Sequencing Tests for Universal Pathogen Detection. Archives of Pathology & Laboratory Medicine 141, 776–786 (2017).

10. Diao, Z., Han, D., Zhang, R. & Li, J. Metagenomics next-generation sequencing tests take the stage in the diagnosis of lower respiratory tract infections. Journal of Advanced Research 38, 201–212 (2022).

11. Rasmussen, T. R., Korsgaard, J., Møller, J. K., Sommer, T. & Kilian, M. Quantitative culture of bronchoalveolar lavage fluid in community-acquired lower respiratory tract infections. Respir Med 95, 885–890 (2001).

12. Gaston, D. C. et al. Evaluation of Metagenomic and Targeted Next-Generation Sequencing Workflows for Detection of Respiratory Pathogens from Bronchoalveolar Lavage Fluid Specimens. J Clin Microbiol 60, e0052622 (2022).

13. Alcolea-Medina, A. et al. Unified metagenomic method for rapid detection of microorganisms in clinical samples. Commun Med 4, 135 (2024).

14. Yin, Y. et al. Enhancing lower respiratory tract infection diagnosis: implementation and clinical assessment of multiplex PCR-based and hybrid capture-based targeted next-generation sequencing. EBioMedicine 107, 105307 (2024).

15. Zhan and Massoumi-Alamouti, S. H. and S. et al. Target capture sequencing of SARS-CoV-2 genomes using the ONETest Coronaviruses Plus. Diagn Microbiol Infect Dis 101, 115508 (2021).

16. Briese, T. et al. Virome Capture Sequencing Enables Sensitive Viral Diagnosis and Comprehensive Virome Analysis. mBio 6, e01491–01415 (2015).

17. Lan, Q. et al. Evaluation of hybrid capture-based targeted and metagenomic next-generation sequencing for pathogenic microorganism detection in infectious keratitis. BMC Infect Dis 25, 1211 (2025).

18. Kok, N. A. et al. Host DNA depletion can increase the sensitivity of Mycobacterium spp. detection through shotgun metagenomics in sputum. Front Microbiol 13, 949328 (2022).

19. Chen, Y., Fan, L.-C., Chai, Y.-H. & Xu, J.-F. Advantages and challenges of metagenomic sequencing for the diagnosis of pulmonary infectious diseases. Clin Respir J 16, 646–656 (2022).

20. Gu, W., Miller, S. & Chiu, C. Y. Clinical Metagenomic Next-Generation Sequencing for Pathogen Detection. Annu Rev Pathol 14, 319–338 (2019).

21. Mamanova, L. et al. Target-enrichment strategies for next-generation sequencing. Nat Methods 7, 111–118 (2010).

22. Chen, W. et al. A case report of confirmed difficult pulmonary tuberculosis based on the hybrid capture-based tNGS method. BMC Pulm Med 25, 64 (2025).

23. Huygens, S. et al. Diagnostic Value of Microbial Cell-free DNA Sequencing for Suspected Invasive Fungal Infections: A Retrospective Multicenter Cohort Study. Open Forum Infect Dis 11, ofae252 (2024).

24. Aerts, R., Feys, S., Mercier, T. & Lagrou, K. Microbiological Diagnosis of Pulmonary Aspergillus Infections. Semin Respir Crit Care Med 45, 21–31 (2024).

25. Rand, K. H. et al. Relationship of Multiplex Molecular Pneumonia Panel Results With Hospital Outcomes and Clinical Variables. Open Forum Infect Dis 8, ofab368 (2021).

26. Kasper, D. L. & Fauci, A. S. Harrison’s Infectious Diseases. (McGraw-Hill Medical, 2010).

27. Langmead, B. & Salzberg, S. L. Fast gapped-read alignment with Bowtie 2. Nat Methods 9, 357–359 (2012).

28. Li, H. et al. The Sequence Alignment/Map format and SAMtools. Bioinformatics 25, 2078–2079 (2009).

29. Wood, D. E., Lu, J. & Langmead, B. Improved metagenomic analysis with Kraken 2. Genome Biol 20, 257 (2019).

30. Camacho, C. et al. BLAST+: architecture and applications. BMC Bioinformatics 10, 421 (2009).

31. Mortazavi, A., Williams, B. A., McCue, K., Schaeffer, L. & Wold, B. Mapping and quantifying mammalian transcriptomes by RNA-Seq. Nat Methods 5, 621–628 (2008).

32. Parks, D. H., Imelfort, M., Skennerton, C. T., Hugenholtz, P. & Tyson, G. W. CheckM: assessing the quality of microbial genomes recovered from isolates, single cells, and metagenomes. Genome Res 25, 1043–1055 (2015).

33. Baker, D. N. & Langmead, B. Dashing: fast and accurate genomic distances with HyperLogLog. Genome Biol 20, 265 (2019).

34. Seemann, T. Prokka: rapid prokaryotic genome annotation. Bioinformatics 30, 2068–2069 (2014).

35. Page, A. J. et al. Roary: rapid large-scale prokaryote pan genome analysis. Bioinformatics 31, 3691–3693 (2015).

36. Katoh, K. & Standley, D. M. MAFFT Multiple Sequence Alignment Software Version 7: Improvements in Performance and Usability. Mol Biol Evol 30, 772–780 (2013).

37. Pertea, G. & Pertea, M. GFF Utilities: GffRead and GffCompare. F1000Res 9, ISCB Comm J-304 (2020).

38. Palmer, J. M. & Stajich, J. Funannotate v1.8.1: Eukaryotic genome annotation. Zenodo 10.5281/zenodo.4054262 (2020).

39. Stanke, M., Steinkamp, R., Waack, S. & Morgenstern, B. AUGUSTUS: a web server for gene finding in eukaryotes. Nucleic Acids Res 32, W309–312 (2004).

40. Emms, D. M. & Kelly, S. OrthoFinder: phylogenetic orthology inference for comparative genomics. Genome Biol 20, 238 (2019).

