## Supplemental Methods for "ONETest PathoGenome: A Multi-Cohort Evaluation of an Optimized NGS Assay for Detection of Lower Respiratory Pathogens in Bronchoalveolar Lavage"

### Methods (Supplementary)

#### Microbiologic Culture Methods

Cultures were performed by standard methods, and all isolates were identified, and antimicrobial susceptibility testing performed by VITEK MS mass spectrometry and by Vitek II (AST GP 72, AST GP 78, AST GN 73, and AST XN06 BioMerieux, Durham, NC, USA). All cultures were plated using the standard quadrant technique onto blood, chocolate, and MacConkey agar (Becton Dickinson, Sparks, MD, USA), incubated in 5% CO2 at 36°C, and examined for growth over the next 24–48 hours, as fully described[25]. If ordered, specimens were run on the PN Panel per the manufacturer's instructions using the FilmArray 2.0 instrument (BioFire Diagnostics, Salt Lake City, UT, USA). Cultures were tested for mycobacteria using standard Lowenstein Jensen media and broth cultures using the BACTEC MGIT 960 held for 42 days. Prior to September 16, 2023, these cultures were performed in the Shands Hospital laboratory, but after that time (N=42) they were sent to a reference laboratory, ARUP, Salt Lake City, UT.

#### OT: DNA extraction

Residual BAL specimens remaining after standard hematology laboratory cell counts from the UF Health Shands Hospital Core Laboratory were frozen at -70° C until processing and used for clinical validation. For each sample processed by OT, only ~700 µL of BAL was available. Due to this constraint in specimen volume, samples did not undergo the normal concentration and processing steps associated with conventional microbiological and mycobacterial culture. For cohort 1, total nucleic acids were extracted from a 500 µL aliquot of frozen excess BAL using the magnetiQ Blood and Cell DNA Extraction Kit and the miQron™ nucleic acid purification instrument (Galenvs, Montreal, Canada). Cohorts 2 and 3 were processed using the MagMAX™ Microbiome Ultra Kit with bead tubes (Thermo Fisher Scientific, Canada) according to the manufacturer's instructions. Extracts were eluted in 50 µL elution buffer and quantified using the Qubit HS DNA assay and NanoDrop One (Thermo Fisher Scientific, Canada).

#### OT: Library preparation, QP enrichment, and NGS

Illumina-compatible TmGS libraries were prepared from total nucleic acids using the OT assay kit according to the manufacturer's instructions (Fusion Genomics Corp., Richmond, BC, Canada). Briefly, nucleic acids were enzymatically fragmented, ligated to indexed Illumina adapters, amplified (18 cycles), and hybridized with biotin-labeled QPs in the presence of adapter-specific blocking reagents for 4 h at 50 °C. Target–probe duplexes were captured using streptavidin-coated magnetic beads and purified by iterative high-stringency washes.

For cohorts 1 and 2, enriched libraries were sequenced by Novogene Co. (Redwood, CA, USA) using 2 × 150 bp paired-end sequencing on Illumina NovaSeq instruments. For cohort 1, prepared libraries were split into two equal aliquots for WmGS and OT. For LoD experiments and cohort 3 (LDT validation), the same workflow was performed using full walk-away automation on a Hamilton STAR liquid-handling system (Fig. S1), followed by sequencing on an Illumina NovaSeq instrument at UF Pathology Laboratories (Rocky Point). Raw sequencing reads (Mean 5.5 million paired-end reads per sample across cohorts 1–3, Table S1) were deposited in the NCBI Sequence Read Archive (SRA) after removal of human-derived reads, under BioProject PRJNA1402314 (Table S11).

**OT: Quantum probe-design pipeline**

The LRT assay was developed by building on our original QP[15] expanding the design to cover species associated with LRTIs and related lineages across the core-genome dataset (Table S10A–B). To improve species- and subspecies-level detection, QPs incorporate frequently observed nucleotide variants (>1%) retrieved from public databases across multiple strains within each taxon MSA. Probe spacing was set at 60 nt to enable recovery of near-complete protein-coding genes with 2X coverage, and probe design ensured coverage of GC-rich regions ranging from 35% to 75%. This design yielded 6.2 million genetic-distance–tolerant, core-genome QPs with an average length of 120 nucleotides (Fig. S3).

**OT: Microbial dataset assembly**

The OT panel of known human pathogens and normal flora was developed through a review of clinical expert consensus guidelines and infectious disease literature[26], supplemented with genome sequence references from high-quality public microbial datasets. These included 661,000 sequences from the European Nucleotide Archive (ENA) and the National Center for Biotechnology Information (NCBI). For QP design and data analytics, we selected a subset of references by excluding atypical genomes and retaining strains annotated as human-related, clinical, or pathogenic. Genomes were further filtered to remove low-quality assemblies and highly redundant sequences. From the remaining genomes, we selected a diversity-maximized subset that preserved representation across known pathogens including obligatory and opportunistic taxa, yielding a curated reference dataset of 47,038 genomes.

**OT: Data analytics – Genome assembly**

We updated our original data analytics pipeline[15] to incorporate the lower-airway microbial community. Reads were quality-filtered as described previously[15] and mapped against the human reference genome (GRCh38.p13, release 35) and the plasmid database COMPASS using Bowtie2[27] v2.4.2, with mapped reads removed. Remaining reads were analyzed in parallel by (i) alignment to the OT curated reference dataset using Bowtie2 (“very-sensitive-local –score-min G,100,9”), with duplicate and high-quality reads identified using Samtools[28] v1.11, and (ii) k-mer–based classification[29] using Kraken2 PlusPF (v2024) with --confidence-level 1. Finally, comparative assemblies were performed to reconstruct consensus genome sequences using Samtools and in-house tools. Poor-quality bases (<Q15), and nucleotide variants were filtered unless they meet the following criteria: (1) quality score of ≥Q15, (2) support from more than one forward-aligned and one reverse-aligned read, and (3) presence in at least 25% of the reads, with a maximum depth of 200,000 allowed during pileup. Taxonomic classification and species’ affiliation of the constructed consensus were further validated using external references using Kraken2 and BLAST+[30]. Finally, all detected species are qualified based on mapped Fragments Per Kilobase Million[31] (FPKM), breadth of coverage (BoC), assembly length, relative abundance, and uniquely mapped reads (MAPQ > 40), derived from BAM files using Samtools. The OT pipeline is implemented in C/C++ and Python, integrating in-house software with third-party tools, as described by Zhan and Alamouti et al. (2021).

**OT: Microbial qualification and categorization (remaining)**

Following genome assembly and species-level validation, the OT pipeline applies a statistical framework that evaluates the entire microbial profile of each sample to distinguish true microbial signal from background baseline and technical noise. This process consists of two key steps: microbial signal qualification and categorization. During signal qualification, OT removes any microbial signal detected in negative controls, including no-template controls (NTCs) when available. The remaining signals are declared detected and then categorized using an outlier-based framework tailored to the OT assay. This framework was calibrated using outlier detection results from 360 clinical BAL samples (cohort 2), allowing empirical, species-specific thresholds to be defined based on the observed distributions of FPKM, relative abundance, assembly length, and uniquely mapped reads. Detected target signals that exceed species-specific upper outlier thresholds across all metrics [FPKM, relative abundance, assembly length/breath of coverage] are classified as strong, those exceeding a lower outlier threshold across at least two metrics are classified as moderate, and all remaining signals that pass negative/NTC filtering but meet the threshold for only one metric are designated as weak. In parallel, any microbial species from these groups that appear in the commensal database are annotated as normal flora. This categorization framework is then used throughout the OT validation studies, where the detection and classification performance (strong, moderate, weak) is benchmarked against conventional diagnostic culture results.

**OT: Core-genome and ortholog identification**

Microbial pathogens were selected from the curated reference dataset to cover 50 phylogenetic families relevant to LRTIs, encompassing 254 bacterial, fungal, and DNA viral species and related lineages (Table S10A). For bacterial core-genome identification, genome quality was assessed using checkM[32] v1.0.13, and assemblies with <90% completeness or >10% contamination were excluded. To reduce redundancy, genomes were clustered at 99% nucleotide identity using Dashing[33]. Genomes were re-annotated using Prokka[34] v1.14.5 (GFF3 format) and analyzed with Roary[35] (min identity 90% BLASTP identity) to define core genes (≥95% of genomes), which were aligned using MAFFT[36] v7.475 (--auto). For viral and fungal ortholog identification, OT applied a parallel workflow. Coding sequences were extracted from existing annotations using gffread[37], while genomes lacking annotations were processed using Funannotate[38] v1.8.16 and Augustus[39] for ab initio gene prediction. Orthologs were inferred using OrthoFinder[40] v2.5.5, with core orthologs defined as genes present in >90% of sequenced isolates (within a species or closely related species, depending on genome availability), and aligned using MAFFT. The final MSA dataset comprised 318,735 bacterial core-genome sequences, 367,764 fungal orthologs, and 907 viral orthologs (Table S10B), which were used for QP design and data analytics within FusionCloud.

**Quality-Not-Sufficient determination**

Sample adequacy and sequencing quality were assessed using an internal QNS metric based on human ribosomal DNA signal. Reads mapping to the human 18S and 28S rDNA genes were used to calculate a composite sample quality index incorporating weighted mean sequencing depth, breadth of coverage, and total read counts across these loci. QNS assessment was applied only to samples in which no microbial targets were detected, to distinguish true negative results from negatives potentially attributable to insufficient sample quality. Samples with a quality index value below a predefined threshold were designated as QNS and excluded from downstream performance analyses. This approach was applied uniformly across all study cohorts.

**Precision/Reproducibility**

Three BAL clinical samples positive for microbial pathogens by both culture and OT were used as positive controls, with concurrent negative controls. For each positive-control BAL sample, two 500 µL aliquots were extracted using the protocol described herein. Extracts (2 × 50 µL) were pooled (100 µL), mixed, divided into five equal aliquots per sample (n = 15), and stored at −20 °C until processing. Aliquots were analyzed across three independent OT runs, including triplicate aliquots per sample within a single run (intra-assay reproducibility) and single aliquots per sample in two additional runs performed on separate days (inter-assay reproducibility). Sequencing quality-control metrics and species detection were evaluated using predefined detection thresholds derived from the BAL training dataset. Successful control performance was defined as no reportable detections above threshold in negative controls and detection of all expected organisms above threshold in positive controls across all runs.

**Continuity measurement/drift control**

Longitudinal assay performance was monitored using a predefined control species (*Listeria monocytogenes*) and human DNA, with a Drift Ratio calculated as the ratio of *L. monocytogenes* FPKM to human FPKM in designated negative-control samples for each sequencing run. Laboratory-defined acceptance limits for the Drift Ratio were established during assay validation based on the distribution of values across multiple runs and are used to flag potential shifts in assay performance over time. Values that remain within the predefined laboratory acceptance criteria, which permit a coefficient variation (CV) of ≤30% based on validation data, are taken to represent stable assay performance.
