## Supplemental Data 1 for "ONETest PathoGenome: A Multi-Cohort Evaluation of an Optimized NGS Assay for Detection of Lower Respiratory Pathogens in Bronchoalveolar Lavage"

### Supplement Figures

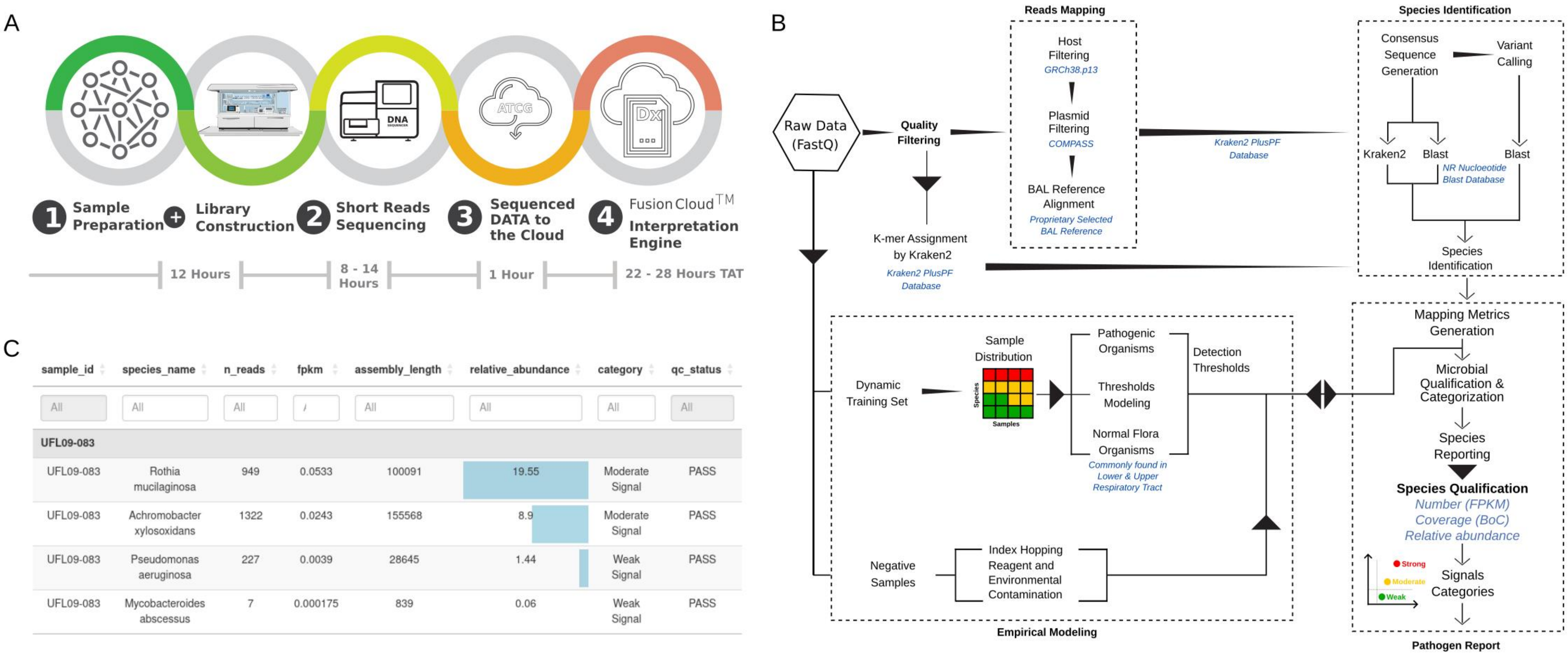


| sample_id | species_name | n_reads | fpkm | assembly_length | relative_abundance | category | qc_status |
|---|---|---|---|---|---|---|---|
| All | All | All | / | All | All | All | All |
| UFL09-083 | | | | | | | |
| UFL09-083 | Rothia mucilaginosa | 949 | 0.0533 | 100091 | 19.55 | Moderate Signal | PASS |
| UFL09-083 | Achromobacter xylosoxidans | 1322 | 0.0243 | 155568 | 8.9 | Moderate Signal | PASS |
| UFL09-083 | Pseudomonas aeruginosa | 227 | 0.0039 | 28645 | 1.44 | Weak Signal | PASS |
| UFL09-083 | Mycobacteroides abscessus | 7 | 0.000175 | 839 | 0.06 | Weak Signal | PASS |

**Figure S1. ONETest™ (OT) end-to-end workflow: (A) ONETest™ laboratory workflow.** Total nucleic acid samples are processed using proprietary Quantum Probes reagents. Captured libraries are transferred to a **Hamilton robotic platform for full hands-off, walk-away processing**, including automated liquid handling, library preparation, sequencing and analysis. The fully automated workflow enables a complete sample-to-report turnaround time of approximately 22–28 hours, encompassing library preparation, sequencing (8–14 hours), data transfer, and analysis. **(B) FusionCloud™ analytical pipeline**. Raw FASTQ files undergo quality-control filtering, followed by parallel k-mer–based taxonomic assignment (Kraken2 with PlusPF database) and read-mapping steps. Species identification is supported by consensus sequence generation, variant calling, and confirmatory BLAST analysis. Quantitative metrics are generated and evaluated against empirically derived detection thresholds. These thresholds are trained using a dynamic reference set incorporating pathogenic organisms, normal respiratory flora distributions, negative/NTC controls, and models for index hopping and reagent contamination, enabling robust microbial qualification and categorization. Signals are categorized as weak, moderate, or strong based on the cumulative number of outlier thresholds exceeded across three key metrics: FPKM, relative abundance, and breadth of coverage. **(C) OT clinical report**. The final report summarizes detected organisms at the species level, including quantitative abundance metrics, categorical signal strength, and quality-control status, providing an interpretable and actionable diagnostic output suitable for routine clinical use.

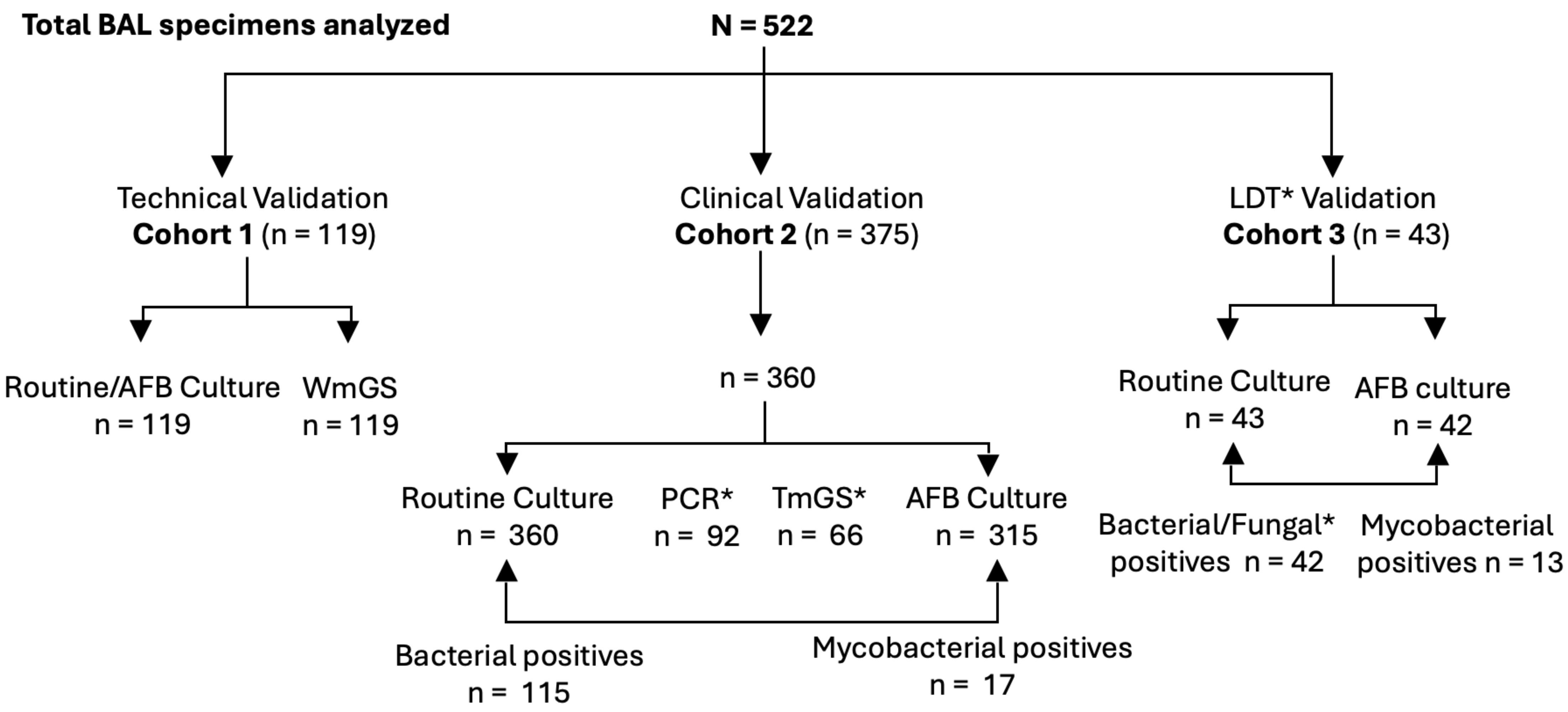


**Figure S2. Total BAL Clinical specimens analyzed by ONETest (OT):** Excess bronchoalveolar lavage (BAL) specimens obtained from the UF Shands Hospital analyzed by OT across three cohorts. QNS exclusion*: Samples that failed internal quality-not-sufficient (QNS) criteria for sample adequacy and sequencing quality, based on human rDNA signal, were excluded; 15 specimens were removed from Cohort 2. OT was benchmarked against culture in all cohorts and against whole-metagenome sequencing (WmGS) in Cohort 1. Acid-fast bacilli (AFB) testing and other repeat bacterial cultures (Table S3) were not mutually exclusive – i.e., the same specimen could undergo multiple tests including subsets of cohort 2 tested by PCR* (BioFire® Pneumonia Panel) and TmGS* (MicroGenDX). LDT* Laboratory development Test validation in a CAP-accredited environment. Bacterial/Fungal*: n=40 when considering bacterial pathogens (i.e., not normal flora) and mycelial fungal spp. growth.

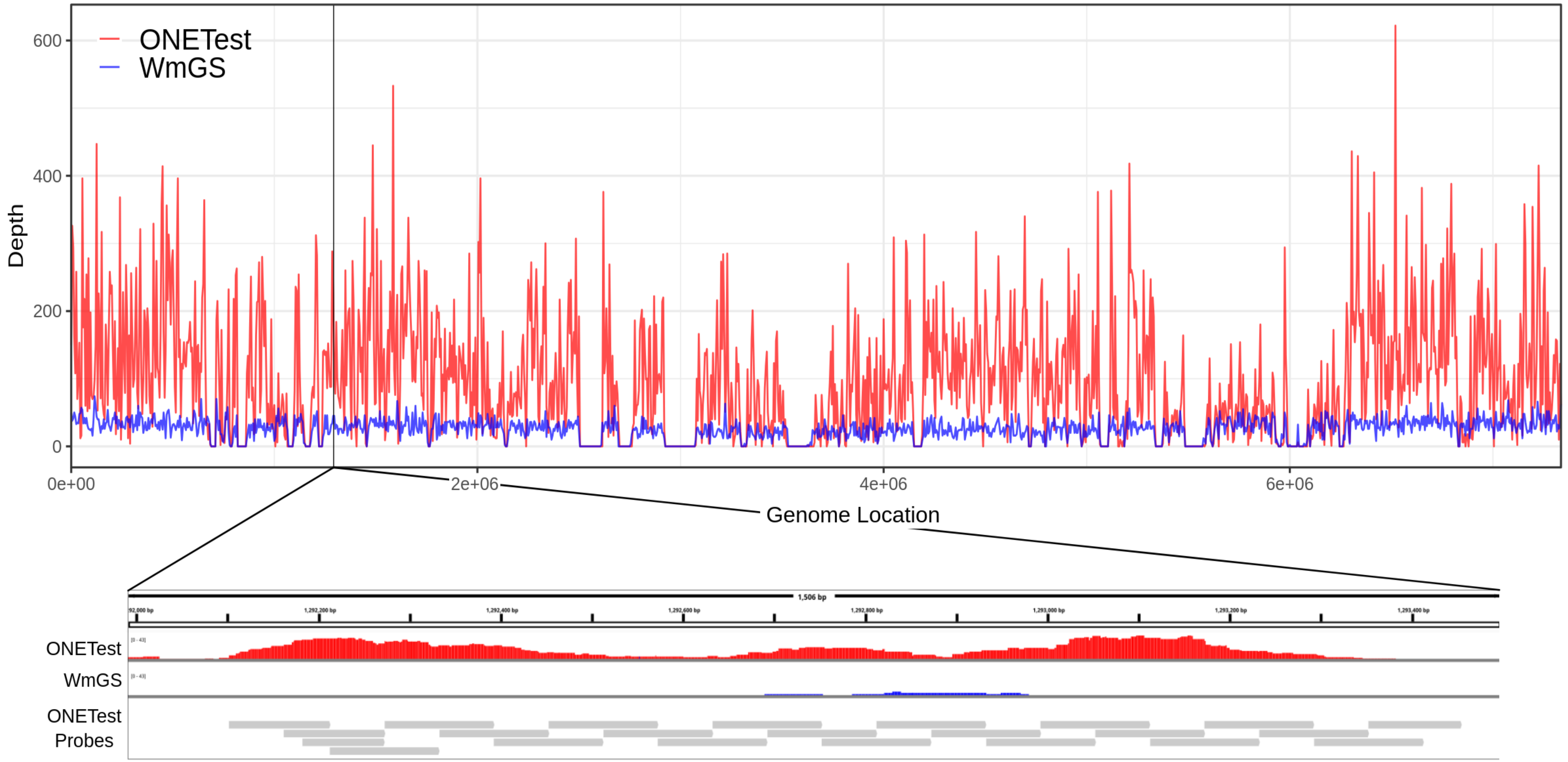


**Figure S3. Core-genome capture of *Pseudomonas aeruginosa* using ONETest Quantum-Probes™**
Genome-wide read depth profiles for ONETest Pathogenome (red) and whole-metagenome shotgun sequencing (WmGS, blue) across the *P. aeruginosa* genome. The ONETest design covers 4,171 core genes with a total of 46,287 core-genome probes, providing at least >2× tiling density across target regions to capture the within-species diversity. The upper panel shows per-base sequencing depth along the full genome, illustrating dense core-genome capture by ONETest compared with lower, relatively uniform coverage by WmGS. The lower panel zooms into a representative core gene highlighting probe tiling probe tiling (grey bars) and the corresponding read-depth profile, demonstrating how high-density, diversity-aware tiling supports continuous, high coverage across the targeted locus.

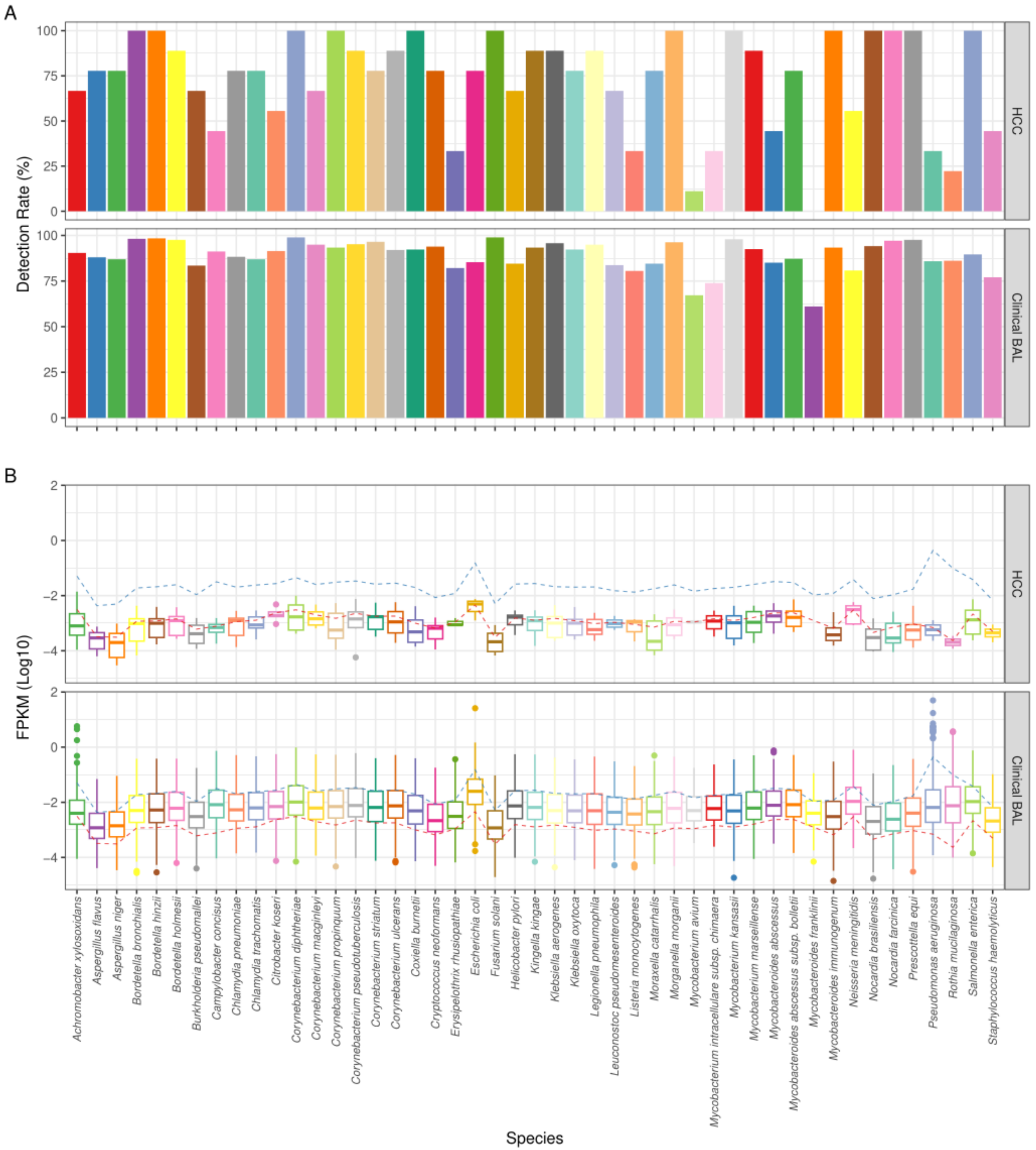


**Figure S4. Overview of contaminants in Bronchoalveolar lavage clinical data:** The detection rate **(A)** and the FPKM **(B)** of the 47 contaminants found in control and human cell controls, and their corresponding values in the 360 clinical BAL samples. The color-dashed lines indicate the FPKM profiles for HCC, and Clinical BAL, represented by red, green, and blue, respectively.

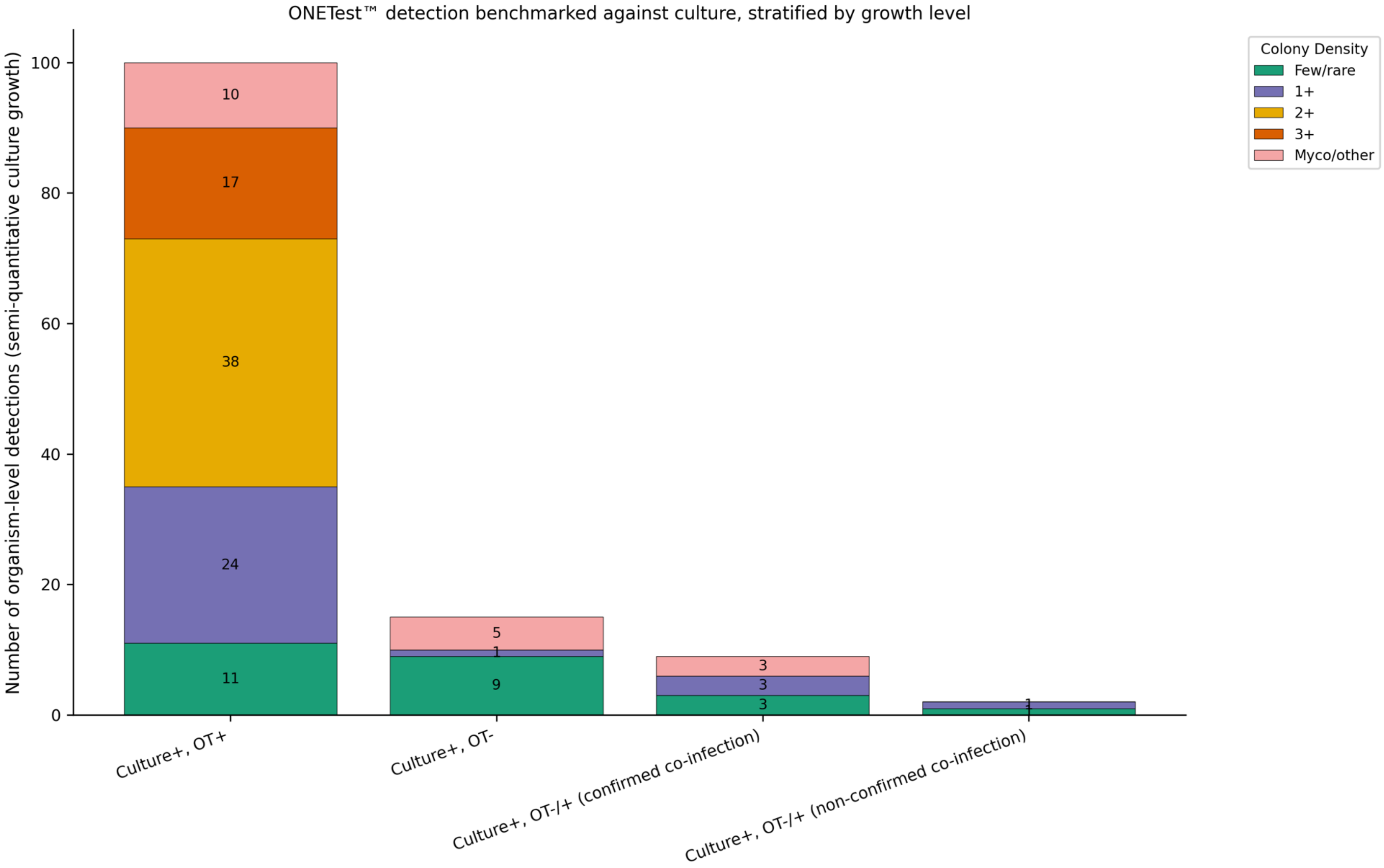


**Figure S5. ONETest™ sample detection benchmarked against culture, stratified by growth level.** Stacked bars show culture-positive organisms from BAL samples classified Culture+, OT+ (TP); Culture+, OT− (FN no co-infection); Culture+, OT−/+ (TP confirmed co-infection), and Culture+, OT−/+ (FN, non-confirmed co-infection). Bar segments indicate semi-quantitative culture colony density: few/rare, 1+, 2+, 3+, Myco/other, representing mycobacterial or non-quantified growth

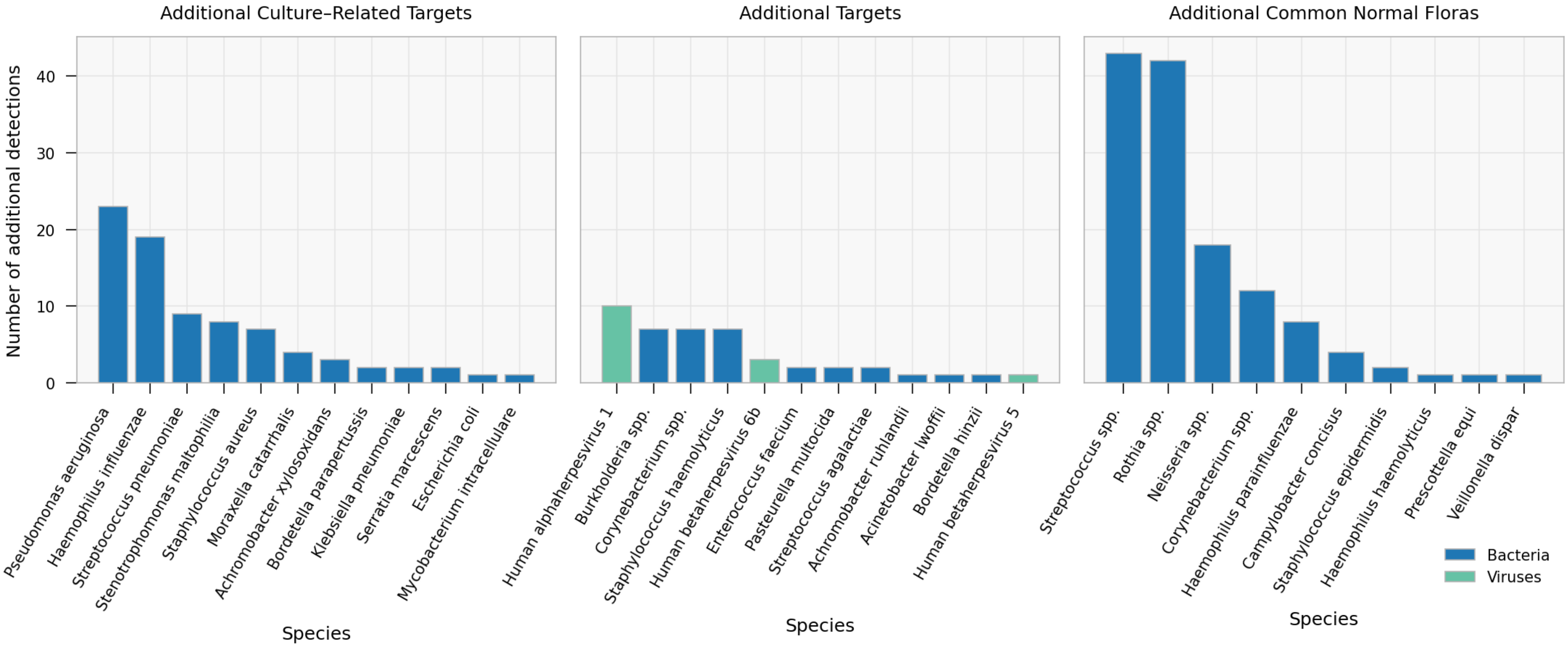


**Figure S6. Composition of additional ONETest™ Pathogenome detections and normal flora by species.** Bar plots summarize species detected by OT that were not primary culture-confirmed pathogens, stratified into three biologically and clinically relevant categories. **Additional culture-recoverable species** (left panel) show OT false-positives corresponding to species that are part of the routine culture menu but were not recovered by culture in the corresponding specimen (81 additional detections across 11 species). **Additional OT-only species** (middle panel) represent OT detections outside the species detected by standard culture workup in cohort 2 (45 additional detections across 21 species). **Additional common normal floras** (right panel) show OT calls interpreted as likely commensal flora in BAL specimens (132 detections across 26 species). Bar height reflects the number of additional detections per species (species-by-sample positive calls). Bars are colored by taxonomic group. Species belonging to the same genus are collapsed to genus-level labels for *Burkholderia*, *Corynebacterium*, *Streptococcus*, and *Neisseria* (full species panel in Table S8).

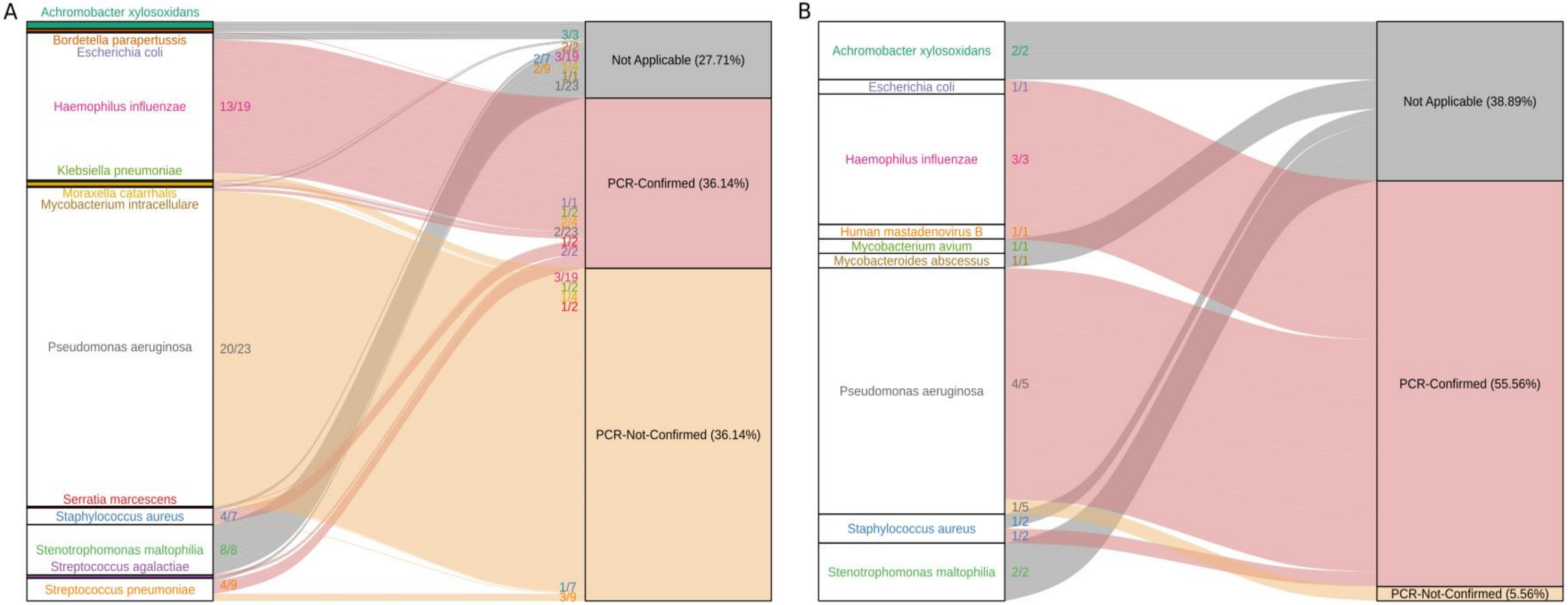


**Figure S7. Distribution of cultured-species ONETest false positives confirmed by PN Panel PCR.** The Sankey charts illustrate the proportion of OT-positive isolates that were not identified through culture but confirmed by PCR (pink) or not confirmed (yellow). "Not applicable" designation when a call was not tested by PCR or when the species is not part of the PN panel list. Chart (**A**) depicts the OT false positive results confirmed by PCR in cohort 2, which includes 81 isolates. Chart (**B**) shows the ONEtest false positive results confirmed by PCR in cohort 3, totaling 16 isolates.

#### Successful Detection of Bacteria from Bronchoalveolar Lavage (BAL) Specimens with Different Concentrations - 0.005, 0.05, 0.5 McFarland

| Sample | Bacteria Mix | BAL Background | Detection Result (FPKM BAL 0.05 to 1.57, BoC 0.06% to 38.39%) | | | | |
|---|---|---|---|---|---|---|---|
| S1 | 2 bacteria spiked | + | Escherichia coli | | Klebsiella pneumoniae | | |
| S2 | | - | Escherichia coli | | Klebsiella pneumoniae | | |
| S3 | 3 bacteria spiked | + | Mycobacterium avium | | Staphylococcus aureus | | Staphylococcus hominis |
| S4 | | - | Mycobacterium avium | | Staphylococcus aureus | | Staphylococcus hominis |
| S5 | 5 bacteria spiked | + | Mycobacterium avium | Escherichia coli | Staphylococcus aureus | Klebsiella pneumoniae | Staphylococcus hominis |
| S6 | | - | Mycobacterium avium | Escherichia coli | Staphylococcus aureus | Klebsiella pneumoniae | Staphylococcus hominis |
| S7 | | + | Mycobacterium avium | Escherichia coli | Staphylococcus aureus | Klebsiella pneumoniae | Staphylococcus hominis |
| S8 | | - | Mycobacterium avium | Escherichia coli | Staphylococcus aureus | Klebsiella pneumoniae | Staphylococcus hominis |

**Figure S8. ONETest detection of Mycobacterium avium in BAL spike-ins with increasing mixture complexity.** Negative BAL matrix (BAL background+) and matched non-BAL controls (BAL background−) were spiked with 2-, 3-, or 5-organism mixtures at three spiked levels (0.5, 0.05, and 0.005 McFarland equivalents) and processed by ONETest. Base concentrations for the 0.5 McFarland suspensions were 3.75×106 CFU/ml for E. coli and *M. avium*; 3.75×105CFU/ml for *Candida* and *S. aureus*; and 3.75×104 CFU/ml for *K. pneumoniae* and *S. hominis*. Subsequent 1:10 and 1:100 dilutions were performed to reach the 0.05 and 0.005 levels, respectively (final titers ranging from 3.75×106 to 3.75×102 CFU/ml). *M. avium* was detected in both 3- and 5-organism mixtures across both backgrounds, indicating retained mycobacterial detectability despite increasing co-spiked organism complexity (FPKM 0.05–1.57; breadth of coverage 0.06%–38%). DNA extraction was performed using MagMax MagMAX™ Microbiome Ultra Kit as described in methods and *Mycobacterium avium a* proteinase K digestion step was added.
